# V-Cornea: A computational model of corneal epithelium homeostasis, injury, and recovery

**DOI:** 10.1101/2025.08.11.669602

**Authors:** Joel Vanin, Michael Getz, Catherine Mahony, Thomas B. Knudsen, James A. Glazier

## Abstract

**Purpose:** To develop a computational model that addresses limitations in current ocular irritation assessment methods, particularly regarding long-term effects, and recovery patterns following chemical exposure or trauma to the cornea.

**Methods:** V-Cornea is an agent-based computer simulation implemented in CompuCell3D that models corneal epithelial homeostasis and injury response. The model incorporates biologically-inspired rules governing cell behaviors (proliferation, differentiation, death) and key signaling pathways including Epidermal Growth Factor (EGF), translating cell-level behaviors to tissue-level outcomes (*in vitro* to *in vivo* extrapolation, IVIVE).

**Results:** V-Cornea successfully reproduces corneal epithelial architecture and maintains tissue homeostasis over extended simulated periods. Following simulated trauma or toxicant exposure, the model accurately predicts healing timeframes of 3-5 days for slight and mild injuries. For moderate injuries with basement membrane disruption, the model demonstrates longer recovery times and emergent dynamic structural disorganization that mimics recurrent corneal erosions, providing mechanistic insights into these pathological conditions.

**Conclusions:** V-Cornea’s modular CompuCell3D implementation is easily extensible to incorporate additional recovery and injury mechanisms. Future versions will include more realistic basement membrane dynamics and explicit representation of stromal tissue and immune response to improve predictivity for moderate injuries. This virtual-tissue approach shows potential not only for toxicological assessments but also for drug discovery and therapy optimization by providing a platform to test interventions and predict outcomes across various injury scenarios.

## 1. Introduction

Toxicological assessment, drug discovery and therapy optimization in the cornea all require the prediction of systemic outcomes from molecular perturbations in the cornea. Accidental exposures to consumer products and industrial chemicals cause eye irritation and corneal damage, creating a significant public health concern. These exposures lead to outcomes ranging from temporary discomfort to permanent vision impairment. We assess ocular irritation potential to ensure the safety of consumer products like shampoos, household cleaning solutions, and laundry detergents, as well as professional products (1). Similarly, developing effective treatments for corneal disorders such as dry eye disease, keratitis, and corneal dystrophies requires understanding how therapeutic compounds interact with corneal tissue at multiple scales. In both toxicology and therapeutics, current approaches often fail to capture the full complexity of molecular interactions and their translation to tissue-level effects, limiting our ability to predict outcomes accurately without extensive animal testing.

A chemical’s inherent properties, exposure conditions, and biological responses influence the severity and persistence of corneal injury following chemical exposure. Assessment of corneal damage found extent of injury to have an important, if not primary, role in eye irritation (2,3). Depth of injury rather than the type of chemical or the mode of chemical damage is particularly significant for recovery time and thus classification of the toxicant. Epithelial injuries typically heal within 3-5 days (mild injury), while injuries affecting the basement membrane and stroma result in prolonged healing times (21+ days) and potential scarring (moderate to severe injury) (4–7).

Toxicologists evaluate corneal damage after toxicant exposure using a variety of experimental methods. Organotypic *ex vivo* assays such as the Bovine Corneal Opacity and Permeability (BCOP) assay and the Isolated Chicken Eye (ICE) test measure opacity, permeability, swelling, and histopathological changes after chemical exposure (8,9). However, ex vivo models have limitations as they lack vascular circulation and can only be maintained for short periods. In addition, interspecies differences reduce their ability to predict human outcomes accurately (10,11). Similarly, *in vitro* assays such as the Reconstructed 3D Human Cornea-Like Epithelium (RhCE) test and the Short Time Exposure (STE) test quantify cytotoxicity and epithelial barrier function but fail to capture the complexity of the *in vivo* cornea, including inflammation responses and wound healing (12,13).

To support in vitro to in vivo extrapolation (IVIVE), we introduce V-Cornea, an agent-based computational model implemented in CompuCell3D (14). V-Cornea’s Virtual Tissue (VT) computer simulations bridge the gap between experimental assays and human outcomes by integrating data from *ex vivo, in vitro*, and cross-species tests to predict human corneal damage and recovery after chemical injury. V-Cornea replicates critical behaviors and interactions of specific tissue components across multiple biological scales, integrating molecular interactions with cellular and tissue-level processes (15,16). When we combine these approaches, our VT simulations complement existing non-animal methods by providing mechanistic insights into how cellular responses in simplified test systems translate to tissue-level outcomes, including the temporal dynamics of injury and repair. While our focus here is on damage and recovery for toxicology, the same model structure can be applied to drug discovery, tissue engineering and therapy optimization (17–22).

### 1.1 Current Approaches in Corneal Modeling

Computer simulations of the cornea typically address distinct biomedical questions following one of three main approaches: (I) finite element analysis (FEA) for understanding mechanical properties, usually representing the cornea as continuous tissue with elastic/viscoelastic properties, focusing on stress-strain relationships without cellular components, (II) mathematical models using partial differential equations (PDEs) to predict wound healing after chemical or abrasive injury, typically represent cell populations as density fields with migration and proliferation rates, and (III) cellular-based models focusing on tissue maintenance.

Researchers using FEA approaches have gained insights into corneal shape changes after refractive surgery, stress distributions in keratoconus, and deformation under varying intraocular pressures (23,24). Over time, adding more realistic material properties to these methods—such as nonlinear elasticity and anisotropic collagen alignment—improved the fidelity of simulated mechanical responses (25). However, corneal FEA-based models define tissues through discretized mesh elements with mechanical constitutive relationships, rather than biological processes governing cell turnover and wound repair. Although some advanced FEA models in other tissues, such as cardiac models (26), do include continuum representations of biological processes like cell turnover and tissue remodeling, to our knowledge, existing corneal FEA models have primarily focused on mechanical properties rather than the biological dynamics of epithelial maintenance and repair that are central to our investigation.

Partial differential equation models address specific aspects of corneal wound healing, such as epithelial cell migration patterns after abrasion, and oxygen distribution during hypoxic conditions (27,28). More recently, mathematical modeling approaches using ordinary differential equations (ODEs) have been applied to understand corneal epithelium maintenance. Moraki et al. (2018.) developed a model based on chemical master equations, but primarily analyzed the derived ODEs to determine constraints on stem cell numbers, division rates, and generation capacity required for maintaining corneal homeostasis. Their model focused on proliferation kinetics and analyzed factors influencing cell populations at steady state, but did not incorporate spatial dynamics or injury response mechanisms. Multicellular approaches investigate how different parameters (replicative lifespan, spatial correlations, and bias) affect the overall tissue homeostasis and pattern formation in corneal basal layer, without explicitly modeling biochemical signaling pathways (30). By simulating individual cells, multicellular models reveal how local cell replication and removal drive tissue-level organization.

Researchers rarely integrate mechanical, chemical, and cellular aspects of corneal biology into a single platform that simulates both normal tissue maintenance and injury response. Our Virtual Cornea (V-Cornea) addresses this gap by blending mechanistic rules for cell behaviors with simplified chemical signaling, compared to actual biochemical pathways but sufficient to capture essential interactions, and tissue architecture, offering a new avenue by creating a comprehensive platform that integrates cellular, mechanical, and biochemical aspects rather than focusing on just one aspect for studying corneal injury and recovery dynamics.

### 1.2 Biological Background

#### 1.2.1 Overview of the Corneal System

The cornea forms the transparent anterior surface of the eye, serving as both a protective barrier and the primary refractive element of the visual system [Fig. 1]. As the outermost tissue, it interfaces directly with the external environment while maintaining optical clarity essential for vision. Structurally, the cornea consists of five distinct layers: the epithelium, Bowman’s layer, stroma, Descemet’s membrane, and endothelium. Each layer contributes uniquely to corneal function, with the epithelium serving as the primary protective barrier against external threats. (31)

**Fig 1.**
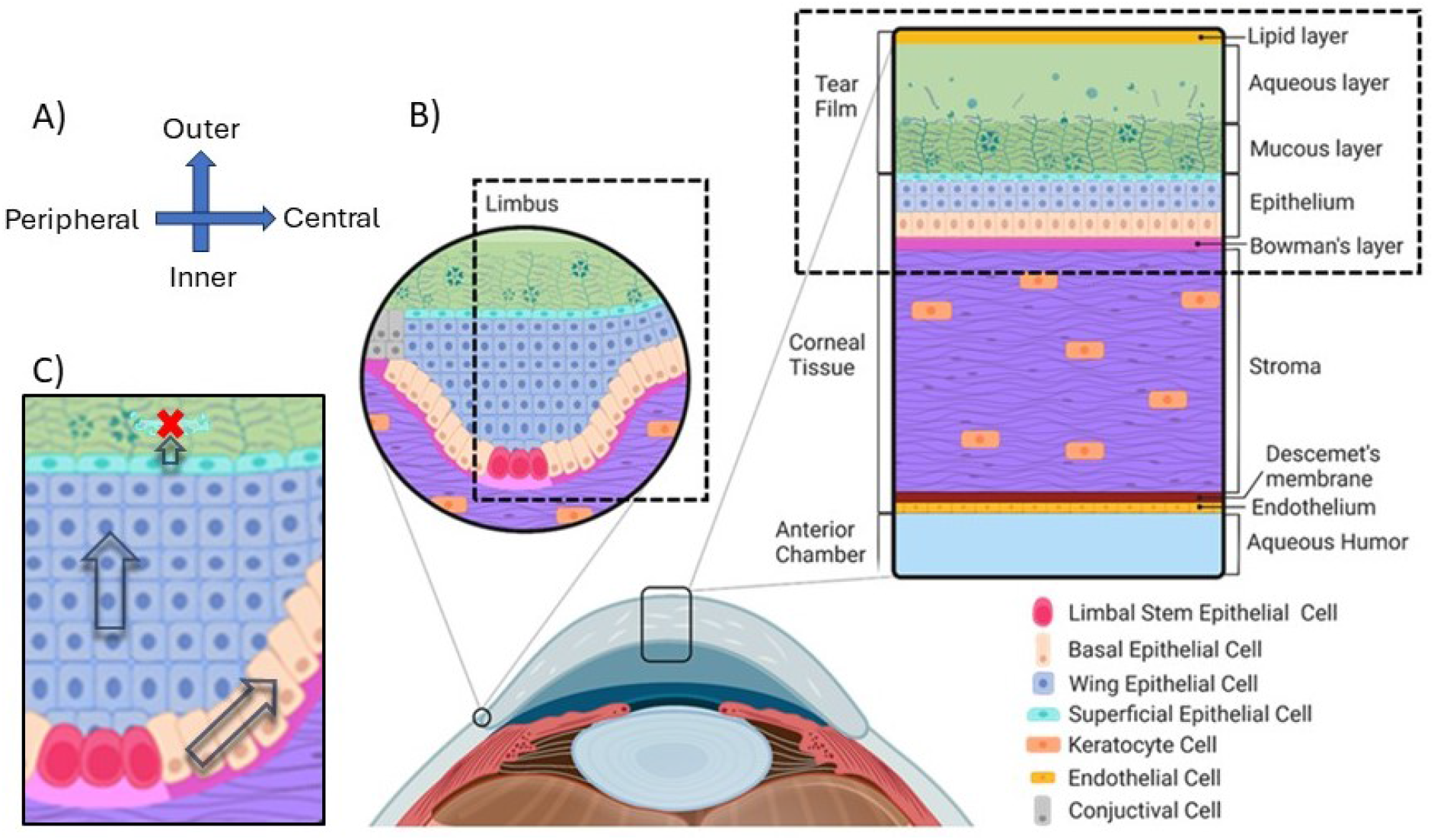
Schematic of Microanatomy proliferation, differentiation, and cell movement during corneal homeostasis. A) In our coordinate system, the x-axis represents the radial position across the cornea, with lower x values corresponding to the Peripheral region and higher x values corresponding to the Central region. The y-axis denotes the depth position, where lower y values indicate the Inner surface (adjacent to the aqueous humor) and higher y values indicate the Outer surface (in contact with the tear film). This orientation allows us to precisely map cellular processes and movements throughout the corneal structure. B) This cross-sectional view shows the epithelium interacting with the tear film on the outer surface and the endothelium contacting the aqueous humor on the inner surface. The limbal region are the source of limbal epithelial stem cells, which are crucial for epithelial renewal. We also show the boundaries of our simulation domain, which focuses on epithelial layer and tear film in the inner limbic region and the peripheral region of the cornea proper. C) We follow the Limbal Epithelial Stem Cell (LESC) hypothesis which describes the cellular dynamics essential for corneal maintenance. LESCs residing in specialized limbal crypts at the corneal periphery divide to produce daughter cells. These daughter cells migrate centripetally toward the central cornea while continuing to divide. As they move inward, they gradually differentiate and shift upward from the basal layer toward the surface, ultimately being shed through desquamation. This continuous process maintains corneal transparency and integrity.

**Fig 2.**
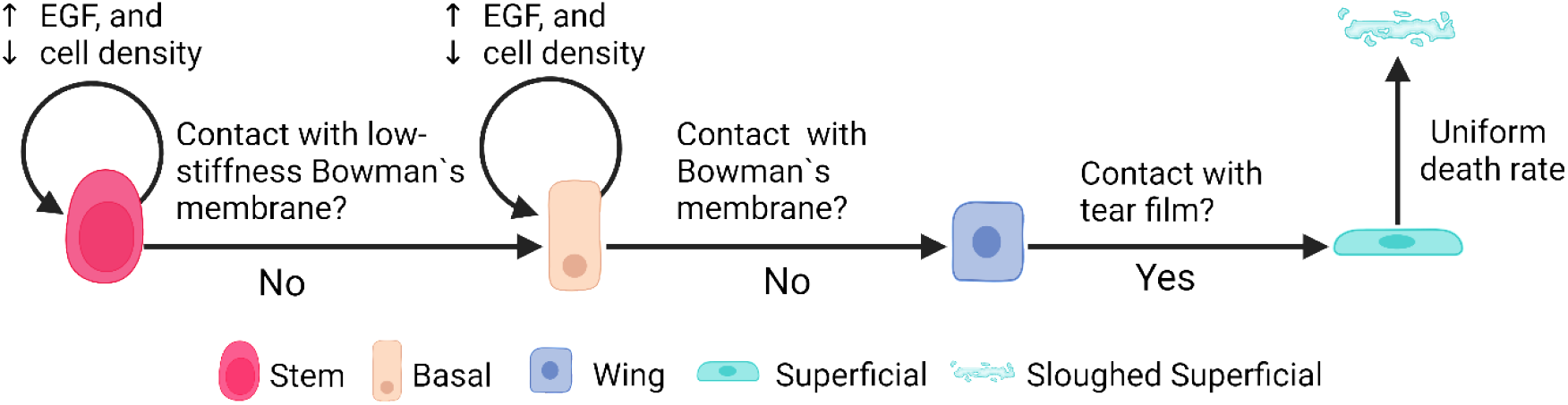
Corneal Epithelial Cell State Transitions and Regulatory Mechanisms. We model corneal epithelial cells progressing through distinct lifecycle states, beginning as stem cells and culminating in their eventual sloughing as superficial cells. We’ve implemented a sophisticated regulatory system where both stem cells (visualized in rose-red) and basal cells (shown in peach-orange) demonstrate proliferation patterns that respond to two key factors: upregulation by Epidermal Growth Factor (EGF) and downregulation by local cell density. We capture critical state transitions based on cellular interactions with their environment. Stem cells transition to a basal state when they lose contact with the lower stiffness regions of Bowman’s membrane (represented in light-pink). These basal cells then transform into wing cells (depicted in blue) upon complete separation from Bowman’s membrane. The final differentiation step occurs when wing cells make contact with the tear film, becoming superficial cells (shown in cyan). To accurately simulate physiological epithelial turnover rates of 7-14 days, we’ve implemented a probability-based sloughing mechanism for superficial cells. This mechanism employs a random draw from a uniform distribution between 0 and 1, compared against one-third of the DaytoMCS value (our conversion factor between simulation time steps and days). This approach ensures we maintain an average transit time of approximately 1.75 days per cell layer, successfully accommodating the varying epithelial thicknesses observed in different regions—from 4 layers at the periphery to 8 layers in the limbal region.

The corneal epithelium constitutes approximately 10% of the total corneal thickness (50-52 μm thick) and forms the outermost cellular layer, situated beneath the tear film. This stratified epithelial tissue comprises 4-8 cell layers that form a tightly regulated barrier (31). The epithelium interfaces with the tear film at its apical surface and with Bowman’s layer at its basal surface. The tear film, with its lipid, aqueous, and mucin layers, both protects the cornea from external insults and delivers growth factors such as EGF that regulate epithelial proliferation and differentiation (32–35).

Between the epithelium and Bowman’s layer lies the epithelial basement membrane (EpBM) (0.05-0.1 μm thick), which serves as both an anchoring structure for epithelial cells and a selective barrier that regulates molecular exchange (36,37). The EpBM plays a crucial role in regulating epithelial-stromal interactions during wound healing by controlling cytokine exchange. When damage disrupts the EpBM, the normal differentiation and adhesion processes are compromised, significantly affecting wound healing outcomes (38–44).

#### 1.2.2 Epithelial Cell Types and Organization

The corneal epithelium is organized into a spatially and functionally distinct hierarchy of cell types, each with specific morphological characteristics and functional roles. These cell populations are maintained through a continuous process of renewal originating from stem cells at the corneal periphery. The epithelium comprises four primary cell types arranged in distinct spatial layers:

##### *Limbal Epithelial Stem Cells* (LESCs)

[Fig. 1 - rose-red cells] reside at the limbus—the border between cornea and sclera—and provide the cellular source for epithelial renewal. These stem cells are characterized by their high nuclear-to-cytoplasmic ratio and slow-cycling nature. LESCs require contact with the limbal basement membrane to maintain their stem cell characteristics (45). They reside within specialized niches in the limbal epithelial crypts, where they receive protection and regulatory signals. These cells retain their proliferative capacity throughout life and serve as the ultimate progenitors for all corneal epithelial cells (46,47).

##### Basal Cells

[Fig. 1 - peach-orange cells] form a single layer of columnar cells that anchor to the basement membrane via hemidesmosomes and retain proliferative capacity. These cells arise from the differentiation of LESCs as they migrate centripetally away from the limbus. Basal cells maintain regular contact with the basement membrane, which is essential for preserving their proliferative potential (48). They typically exhibit a polygonal morphology when viewed from the apical surface and feature prominent cell-cell junctions. Basal cells actively participate in both normal epithelial renewal and wound healing responses, with their adhesion to the basement membrane playing a critical role in epithelial integrity.

##### Wing Cells

[Fig. 1 - blue cells] occupy the intermediate layers (2-3 layers) of the epithelium and derive their name from their characteristic wing-like appearance in cross-section. They arise from the vertical migration and differentiation of basal cells. Wing cells feature a more flattened morphology than basal cells but are not yet fully differentiated. These cells serve as transitional elements that facilitate basal cell movement toward the corneal surface while maintaining epithelial structural integrity (49). Though not proliferative under normal conditions, wing cells retain some capacity to respond to injury signals.

##### Superficial Cells

[Fig. 1 - cyan cells] form the outermost 1-2 layers of the epithelium and represent the terminal stage of epithelial differentiation. These squamous cells are characterized by their remarkably flattened morphology and specialized apical microvilli, which help stabilize the tear film. The most distinctive feature of superficial cells is their formation of tight junctions with a restrictive pore radius of approximately 4 Å (50). These junctions create a highly selective barrier that regulates fluid and molecular exchange between the tear film and underlying corneal tissue. Superficial cells exhibit a finite lifespan of approximately 7-10 days before undergoing a specialized form of non-classic apoptosis and desquamation from the corneal surface (41,51,52).

#### 1.2.3 Epithelial Cell Dynamics and Behaviors

The corneal epithelium maintains its structural integrity and functional capabilities through tightly coordinated cellular behaviors, including proliferation, differentiation, migration, and regulated cell death. These dynamic processes work in concert to ensure continuous renewal while preserving the epithelium’s barrier function.

##### Proliferation and Growth Regulation

Epithelial renewal begins with the proliferation of limbal epithelial stem cells, which exhibit a specialized centripetal division pattern (53). This directional division produces progenitor cells that migrate toward the central cornea, a process crucial for replenishing the central epithelium and preserving corneal transparency (53). LESC proliferation is precisely regulated by intrinsic genetic programs and extrinsic signaling from the surrounding niche environment (47).

Basal cells also contribute significantly to epithelial renewal through their proliferative capacity. Unlike the directed division pattern of LESCs, basal cells follow a random cleavage plane orientation during division. This randomized pattern serves a crucial role in maintaining tissue homeostasis by promoting even cell distribution (48).

Growth factor signaling plays a critical role in regulating epithelial proliferation, with Epidermal Growth Factor (EGF) serving as a primary mitogenic signal (54). EGF stimulates corneal epithelial proliferation particularly during wound healing, while its effect on intact epithelium remains minimal. Interestingly, even with continued EGF treatment, any hyperplastic response eventually returns to normal epithelial thickness, demonstrating the presence of intrinsic regulatory mechanisms (55). The tight junctions formed by superficial cells help regulate EGF penetration into the epithelium, with their restrictive pore radius (approximately 4 Å) (50) limiting the diffusion of larger EGF molecules (~6 kDa) (56) under normal conditions.

##### Differentiation

Epithelial differentiation follows a precisely orchestrated sequence that begins with LESCs and culminates in terminally differentiated superficial cells. This process is primarily regulated by spatial cues and cell-matrix interactions. LESC differentiation into basal cells is triggered when stem cells lose contact with the specialized limbal basement membrane (45,57). This spatial signal initiates the first step in the differentiation cascade as cells migrate centripetally.

Basal cells differentiate into wing cells when they detach from the basement membrane and move vertically within the epithelium. This transition involves significant cytoskeletal reorganization and changes in cell-cell adhesion properties (48).

Wing cells undergo terminal differentiation into superficial cells as they approach the epithelial surface. This final differentiation step involves dramatic morphological changes, including flattening of the cell shape, formation of specialized apical structures, and assembly of tight junctional complexes (49,58).

Throughout this differentiation sequence, cells undergo progressive gene expression changes that reflect their specialized functions at each stage. The entire renewal process, from stem cell division to superficial cell desquamation, takes approximately 7-14 days (59,60), establishing a continuous cycle that maintains epithelial homeostasis.

##### Migration

Cell migration in the corneal epithelium follows distinct patterns that contribute to tissue maintenance and repair. Centripetal migration – following division, LESCs` LESCs and their progeny migrate centripetally from the limbus toward the central cornea. This directional movement establishes a continuous flow of new cells to replace those shed from the central epithelium. This migration is regulated by a complex network of adhesion molecules and extracellular matrix interactions. Specifically, proteins such as laminin and fibronectin provide guidance cues and adhesion substrates that facilitate this directed cell movement (61,62).

Vertical migration – as cells differentiate, they simultaneously migrate vertically from the basal layer toward the corneal surface. This upward movement is facilitated by a combination of cell proliferation in the basal layer, which provides physical impetus, and changes in cellular adhesion properties during differentiation (58).

Wound healing migration – following epithelial injury, basal cells serve as the primary cells that migrate into the wound area (63). This directed movement originate from increased cell proliferation in the limbal region and altered adhesion dynamics, with mechanical constraints from neighboring cells helping create movement toward the wound site (64,65).

##### Cell Death and Shedding

Superficial cells undergo a specialized form of programmed cell death before being shed from the corneal surface. This process differs from classical apoptosis and is designed to maintain barrier integrity even as individual cells are removed from the epithelium (52). The controlled desquamation of superficial cells completes the epithelial renewal cycle and is finely balanced with the production of new cells from the basal layers. This balance ensures that the epithelium maintains a consistent thickness and structural organization while continuously renewing its cellular components.

#### 1.2.4 Epithelial Response to Injury

Following injury, tear production increases as part of the physiological response (66). Damage to the superficial cell layer allows increased influx of growth factors from the tear film, triggering repair responses (5,66,67). Injury-induced breaches in the epithelial barrier allow increased EGF diffusion into the tissue, triggering basal cells to migrate toward the wound site. The healing process progresses through a sequence of basal cell coverage over the wound bed, followed by cell differentiation to restore proper epithelial stratification (63). The severity and nature of the injury, particularly whether it affects the basement membrane, significantly influence the healing outcome (43,68–70). When the basement membrane remains intact, epithelial healing typically proceeds rapidly (3-5 days) and without complications (7,71). However, when the basement membrane is damaged, healing may be delayed, and the risk of recurrent erosions increases due to impaired epithelial adhesion (72,73).

### 1.3 Scope of the V-Cornea Model

Our current implementation of V-Cornea focuses specifically on the epithelium, tear film, and a simplified representation of the basement membrane, as these structures are directly involved in the initial response to chemical exposure and mild to moderate injuries (2,3,6). While the stroma is minimally represented in our model as a spatial constraint, we have not implemented its complex cellular dynamics or collagen organization in this version. (74–76)

We have intentionally excluded detailed representations of the deeper corneal structures (Descemet’s membrane and endothelium (77,78) in this initial model, as they typically become involved only in severe injury scenarios. Additionally, we’ve simplified the immune response aspects of corneal wound healing to focus first on the fundamental epithelial regeneration processes (79,80). These simplifications allow us to create a computationally tractable model that accurately captures the essential dynamics of epithelial homeostasis and recovery from mild to moderate injuries, which are the most common outcomes in accidental chemical exposures. Future iterations of V-Cornea will incorporate these additional structures and processes as we extend the model to simulate more severe injury scenarios.

In this work, we introduce the Virtual Cornea (V-Cornea), a two-dimensional agent-based model (ABM) to elucidate key mechanisms underlying corneal epithelial homeostasis and injury response. While organotypic assays and RhCE tests measure cell death and barrier disruption at endpoints, they fail to capture longitudinal dynamic interactions between cell behaviors and tissue-level organization during injury and recovery. We address this gap by implementing biologically-inspired rules for cell proliferation, differentiation, death and migration to predict how cell-level responses create tissue-level outcomes. Our mechanistic approach reveals how initial cell damage patterns evolve following different recovery scenarios, including behavior mimicking recurrent corneal erosions following basement-membrane disruption. We can simulate extended time periods (up to 6 months) to investigate long-term tissue responses that researchers cannot practically study ex vivo, making our model particularly valuable for understanding how different initial cellular injury patterns lead to distinct adverse outcomes.

## 2. Methods and Implementation

### 2.1 Modeling Approach

We implemented V-Cornea using agent-based modeling (ABM) to represent the spatiotemporal dynamics of corneal epithelial cells. This approach allows individual cells to act as autonomous agents governed by programmed rules, interacting with neighboring cells and their microenvironment. We selected ABM because it effectively captures the coordinated cellular behaviors essential to corneal epithelial homeostasis and wound healing, which are difficult to model using continuum methods.

Our implementation uses CompuCell3D (version 4.6.0), an open-source, cross-platform environment designed for simulating multi-cellular dynamics (https://compucell3d.org) (14). CompuCell3D employs the Glazier-Graner-Hogeweg (GGH) formalism (81), which represents cells as collections of lattice sites and evolves them according to effective energy minimization principles. The ABM approach has demonstrated success in modeling diverse multicellular processes (82) and enables us to represent key cellular mechanisms and the spatially-resolved chemical environment critical for our simulation.

### 2.2 Simulation Domain and Spatial Framework

Our model represents a two-dimensional (2D) radial sagittal section of the corneal limbus and peripheral cornea. The simulation domain is implemented as a discrete square lattice of 200×90 voxels [Fig. 3], where each voxel corresponds to an area of 4 µm^2^, resulting in a total simulated region of 400×180 µm. This spatial scaling allows direct correlation between simulation measurements and biological data. The model operates at a temporal resolution where one Monte Carlo Step (MCS) represents 6 minutes of biological time.

**Fig 3.**
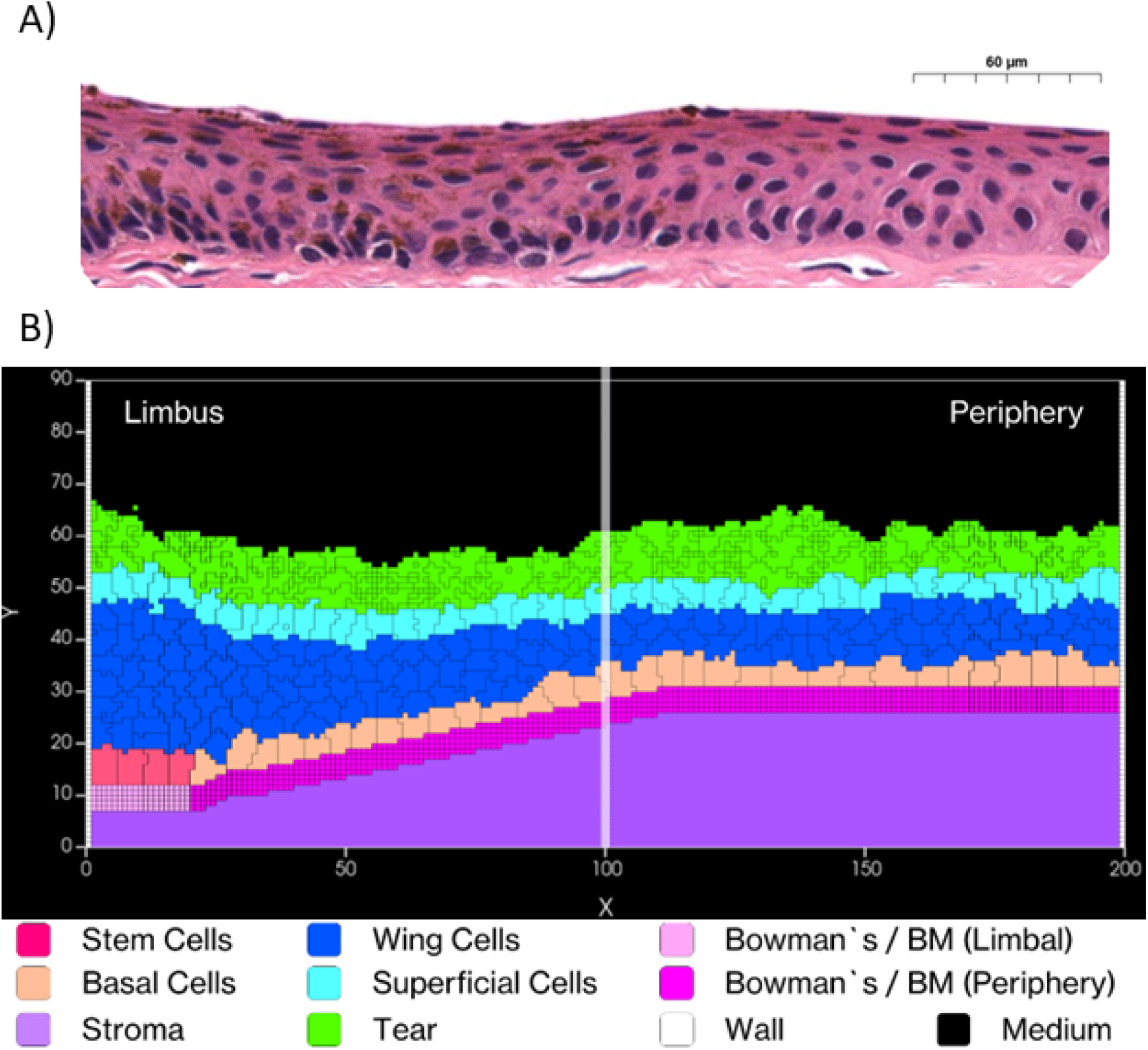
Simulated Corneal Epithelium. A) Histological cross-section of the cornea showing the layered structure. (Image cropped from Sorenson & Brelje, Atlas of Human Histology, 3rd Edition, 2014). Copyright 2014 T. Clark Brelje and Robert L. Sorenson. Used for scholarly and educational purposes under the Fair Use doctrine. B) V-Cornea rendered in CompuCell3D Player. From the bottom, the corneal layers include: the stroma (Lilac-Purple), a merged representation of Bowman’s layer and the epithelial and basement membrane (with low-stiffness limbal regions to the left (Pink) and high-stiffness central regions to the right (Magenta)). Above the basement membrane there is a layer of limbal epithelial stem cells to the left (LESC) (Rose-Red) and basal cells to the right (Peach-Orange). Above the stem cells and basal cells are multiple layers of wing cells (Blue) layer, a single layer of superficial cells (Cyan) and tear-film (Bright Green). The medium (Black) represents the air above the cornea.

We limited our simulation to this region rather than modeling the entire cornea for computational efficiency and because it adequately demonstrates the key cellular dynamics we aim to study. Our domain incorporates anatomically accurate dimensions with an epithelial thickness of 50-52 μm comprising 4-8 cell layers. The limbal region spans approximately 80 μm in width, while the peripheral corneal region extends to approximately 320 μm. We modeled epithelial cell diameters ranging from 10-30 μm (83) and implemented a simulated tear film thickness of 5-10 μm to reflect physiological conditions.

### 2.3 Cellular Components and Agent Representation

#### 2.3.1 Cell Types

V-Cornea implements four distinct cellular agents representing the corneal epithelial cell types, each with specific properties and behaviors. *Limbal Epithelial Stem Cells* (LESCs) [Fig. 4 - rose-red agents] are implemented as proliferative agents that maintain contact with the limbal basement membrane and divide to produce progenitor cells that migrate centripetally (45–47,57). *Basal Cells* [Fig. 4 - peach-orange] are modeled as proliferative cells that maintain contact with the basement membrane, with division following a random cleavage plane orientation to ensure balanced tissue distribution (48). *Wing Cells* [Fig. 4 - blue] function as non-proliferative transitional components in the epithelium (49). *Superficial Cells* [Fig. 4 - cyan] are programmed to form a protective barrier at the apical surface with a finite lifespan, after which they undergo a probabilistic sloughing mechanism (41,51,52).

**Fig 4.**
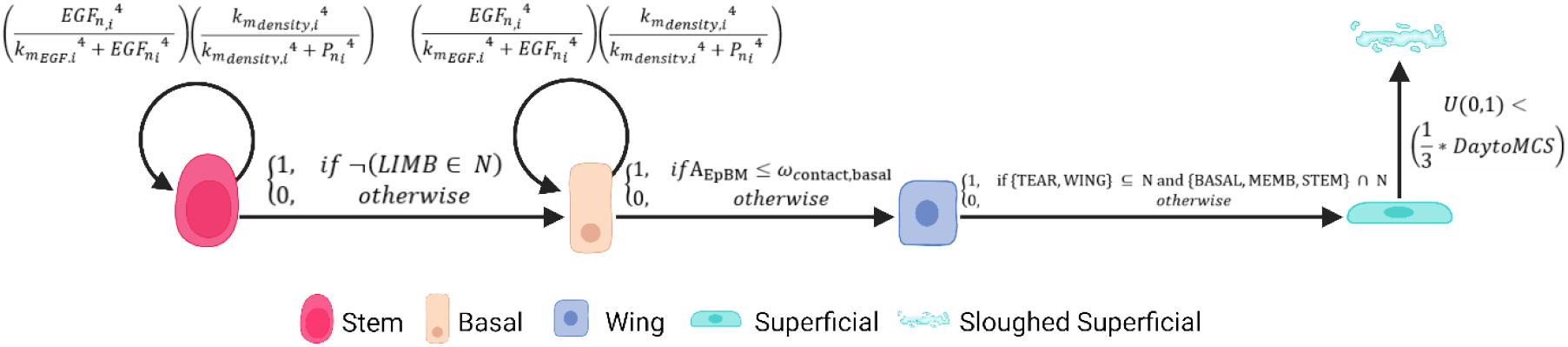
Corneal Epithelial Cell State Transitions and Regulatory Mechanisms. We model corneal epithelial cells progressing through distinct lifecycle states, beginning as stem cells and culminating in their eventual sloughing as superficial cells. We’ve implemented a sophisticated regulatory system where both stem cells (visualized in rose-red) and basal cells (shown in peach-orange) demonstrate proliferation patterns that respond to two key factors: upregulation by Epidermal Growth Factor (EGF) and downregulation by local cell density. We capture critical state transitions based on cellular interactions with their environment. Stem cells transition to a basal state when they lose contact with the lower stiffness regions of Bowman’s membrane (represented in light-pink). These basal cells then transform into wing cells (depicted in blue) upon complete separation from Bowman’s membrane. The final differentiation step occurs when wing cells make contact with the tear film, becoming superficial cells (shown in cyan). To accurately simulate physiological epithelial turnover rates of 7-14 days, we’ve implemented a probability-based sloughing mechanism for superficial cells. This mechanism employs a random draw from a uniform distribution between 0 and 1, compared against one-third of the DaytoMCS value (our conversion factor between simulation time steps and days). This approach ensures we maintain an average transit time of approximately 1.75 days per cell layer, successfully accommodating the varying epithelial thicknesses observed in different regions—from 4 layers at the periphery to 8 layers in the limbal region.

#### 2.3.2 Tear and Epithelial Basement Membrane as Agents

Our model incorporates several environmental components critical for epithelial function. The *Tear Film* [Fig. 3 - bright green] in the simulation, tear film has a fixed level of Epidermal Growth Factor (EGF) which means that EGF is available at all positions of the corneal surface (34,35,54,66). The *Epithelial Basement Membrane* (EpBM) [Fig. 3 - pink/magenta horizontal structure] is modeled as a unified computational entity that combines the biological epithelial basement membrane and Bowman’s layer, represented with two distinct regions: low-stiffness limbal regions (pink) and high-stiffness central regions (magenta)(36,37,39,40,45). The *Stroma* [Fig. 3 - purple] is implemented as a simplified representation as a solid material that exclude the entry of other cells following injury.

### 2.4 Cellular Behaviors and Rules

We implemented cellular behaviors through a set of programmed rules that govern cell differentiation, proliferation, movement, and death. [Table S1–S5]

#### 2.4.1 Growth Dynamics

Cell growth is calculated based on two primary factors: local EGF concentration and mechanical constraints from neighboring cells. Each cell’s growth rate is determined by how its target volume progresses over time. The EGF-dependent growth factor is calculated using a Hill function:

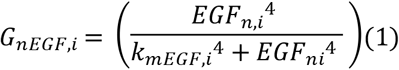

where *EGF*_*ni*_ is the average EGF concentration in cell *i* at time *n*.

Density-dependent growth inhibition is represented by another Hill function:

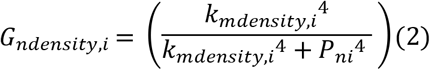

where *P*_*ni*_ is the effective “pressure” inside cell *i* at time *n*, derived from its volume deviation. The total growth rate combines these factors as

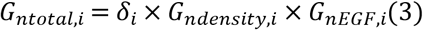

where *δ*_*i*_ is the cell’s intrinsic maximal doubling rate. The parameter values are calibrated to ensure physiologically relevant growth patterns, with detailed formulations provided in Supplementary Information S1.

#### 2.4.2 Mitosis and Cell Division

Cell division is implemented using a volume-based regulation system. We initialize both LESCs and basal cells with a volume of 25 pixels (100 µm^2^) and trigger mitosis when cells reach double their initial volume—50 pixels (200 µm^2^). During division, the parent cell’s volume is equally distributed between the two resulting cells. We implemented different division patterns for different cell types. For LESCs, we oriented the division plane to direct daughter cells centripetally, following a pattern that maintains stem cell populations while producing committed progenitors (61,62,64). For basal cells, we implemented a random cleavage plane orientation to promote even tissue distribution throughout the epithelium (48).

#### 2.4.3 Differentiation

Cell differentiation is implemented as a rule-based process triggered by spatial relationships. LESC to Basal transition occurs when an LESC loses contact with the limbal EpBM [Fig. 4 - first rule] (45,57), represented as

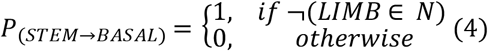

where *N* is the set of neighboring cell types.

Basal to Wing transition is triggered when contact with the basement membrane diminishes (48) below a defined threshold:

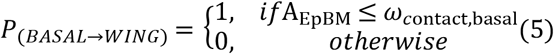

where *A*_*EpBM*_ is the number of pixels in contact with the basement membrane, and *ω*_*contact,basal*_ = 5 pixels are the minimal contact area required to maintain basal phenotype.

Wing to Superficial transition occurs when wing cells reach the uppermost region (49,58) and contact the tear film:

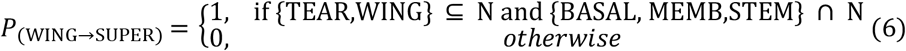

These transitions are deterministic and occur immediately once the condition is met.

#### 2.4.4 Movement

Cell movement is implemented through the Cellular Potts Model framework, where cell positions evolve to minimize the system’s Hamiltonian:

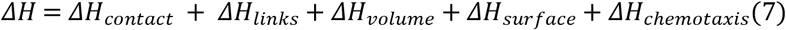

The contact energy term controls adhesion between different cell types, while volume and surface constraint terms maintain cell size and shape. The chemotaxis term enables directed movement in response to chemical gradients. For wound healing scenarios, we implemented directed migration by adjusting chemotactic responses, particularly for basal cells, to move toward the wound site following the basal cell migration hypothesis (63). Cell positions update through the Monte Carlo algorithm that accepts configuration changes when they decrease the system energy or with a Boltzmann probability when they increase it. (Supplementary Information S3)

#### 2.4.5 Cell Death and Sloughing

We implemented two distinct mechanisms for cell removal. Natural sloughing occurs for superficial cells in contact with the tear film through a probabilistic approach:

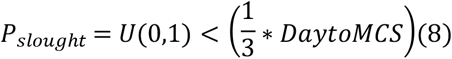

where *U*(0,1) is a uniform random sample in [0,1], and *DaytoMCS* is the conversion factor between days and Monte Carlo Steps (240). This calibrated probability ensures physiological epithelial turnover rates of 7-14 days (84) with an average transit time of approximately 1.75 days per cell layer, accommodating the natural variation in epithelial thickness from 4 layers at the periphery to 8 layers in the limbal region.

For injury-induced death, we model tissue injury through both ablation and chemical exposure mechanisms. Ablation simulates physical trauma by removing cells within a defined circular region and replacing them with tear film to reflect the physiological response of increased tear production following injury. Chemical exposure uses a reaction-diffusion model to simulate toxicant spread, implementing both localized exposures through droplet-like gaussian distributions and broader uniform exposure patterns. When cells encounter chemical concentrations above defined thresholds, we trigger programmed cell death through volume reduction.

Both mechanisms generate distinct spatial patterns of injury but differ in implementation— ablation applies instantaneous geometric removal, while chemical injury incorporates spatiotemporal dynamics dependent on diffusion patterns and concentration gradients. We’ve designed our framework with the flexibility to accommodate more complex chemical behaviors in future iterations. The system can incorporate parameters for chemical reactivity, cellular selectivity, and tear film interactions when experimental calibration data becomes available. This extensibility will enable simulation of diverse chemical exposure scenarios, from caustic agents with direct membrane disruption to surfactants that gradually compromise barrier function. Our current implementation serves as a proof-of-concept demonstration of our ability to distinguish between injury classifications based on depth and recovery dynamics, rather than as a direct simulation of specific chemical mechanisms. We provide the technical specifics of this implementation in Supplementary Information S4.

### 2.5 Tear Film and Epidermal Growth Factor (EGF) Dynamics

We model EGF as a critical signaling molecule using a reaction-diffusion system with two key parameters: diffusion coefficient and decay rate. While our model explicitly implements EGF, this signaling pathway effectively represents an abstraction of multiple growth factors and signaling mechanisms involved in corneal homeostasis and wound healing, including neural regulation (85), inflammatory mediators (86), and other epithelial growth factors (4). This simplified representation allows us to capture essential regulatory dynamics without the computational complexity of modeling each signaling pathway individually. The tear film serves as a constant EGF source at the epithelial surface. The EGF concentration field *c*_*EGF*_(*x,y,t*) evolves according to the equation:

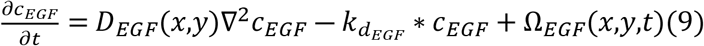

where *D*_*EGF*_(*x,y*) is the cell-type dependent diffusion coefficient, 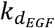 is the global decay rate, and, Ω_*EGF*_(*x,y,t*) represents the net source/sink terms.

We implement cell-type dependent diffusion rates to reflect the distinct permeability characteristics of each epithelial layer. The diffusion coefficient varies by cell type:

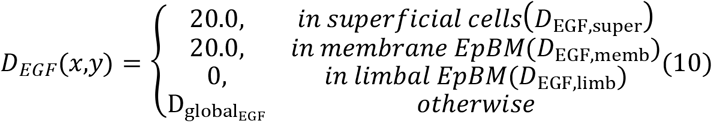

This variation is biologically significant, as the superficial cell layer creates a highly selective barrier through tight junctions. This barrier property is critical for our injury simulations, as damage to the superficial layer allows increased EGF diffusion to basal and stem cells, triggering proliferation and migration responses essential for wound healing.

Our model incorporates varying chemical diffusion rates across different tissue compartments, with 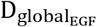 set at 186 voxels^2^/MCS based on experimental data from GelMA hydrogel experiments (87) adapted to our simulation parameters. The global decay constant for EGF 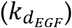 is set to 0.5 per MCS, intentionally higher than the physiological baseline values calculated from EGF’s circulating half-life of 42-114 minutes (88). We selected this elevated decay rate to implicitly incorporate additional processes not explicitly modeled, including cellular uptake, receptor-mediated endocytosis, and sequestration by extracellular matrix components.

We establish boundary conditions to create a physically confined system where EGF cannot escape through lateral boundaries (implemented through wall cells and no-flux conditions), top and bottom boundaries maintain zero concentration, and effective transport is confined to the region between wall cells. This implementation enables us to simulate the dynamic EGF concentration gradients that drive cell behavior during both homeostasis and following injury. This implementation enables us to simulate the dynamic EGF concentration gradients that drive cell behavior during both homeostasis and injury response. As shown in Figure 5, we capture how EGF distribution patterns change dramatically following injury, creating localized regions of elevated concentration at wound sites that trigger cellular responses, and how these patterns normalize as the epithelial barrier is restored during healing.

**Fig 5.**
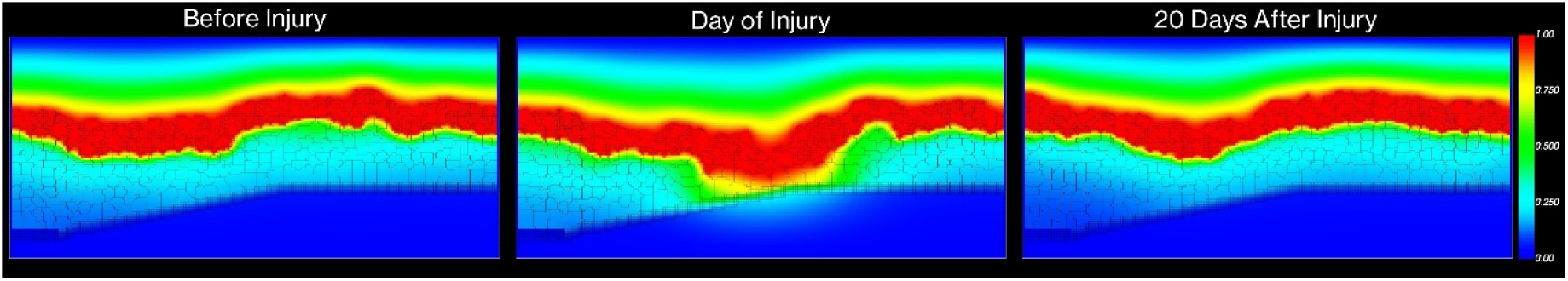
Spatiotemporal Evolution of the EGF Concentration Field During Corneal Injury and Recovery. We capture EGF distribution patterns across three key phases of corneal response to injury in our simulation. In Panel A, we show the baseline EGF gradient under normal physiological conditions, where the tear film maintains a steady-state concentration (normalized to 1.0) that gradually diminishes across the epithelial layers. Panel B reveals the dramatic changes in EGF distribution immediately following injury, with increased EGF influx through the compromised epithelial barrier creating localized regions of elevated concentration at the wound site. In Panel C, we demonstrate the system’s return to homeostasis after 20 days, where the re-established epithelial barrier has restored normal EGF diffusion patterns. We represent normalized EGF concentration through color intensity, with warmer colors (red) indicating higher concentrations and cooler colors (blue) showing lower concentrations. This temporal sequence illustrates how we capture both the acute response to injury and the gradual restoration of normal chemical signaling patterns during wound healing.

## 3. Results

### 3.1 Emergent Behavior and Tissue Homeostasis

V-Cornea demonstrates emergent behavior as complex corneal epithelial structures develop from basic cellular rules. Figure 6 illustrates the day-by-day progression of corneal epithelium generation over 15 simulated days. Starting with only LESCs, tear film, and EpBM, the simulation shows basal cells proliferating and migrating centripetally toward the central cornea during the first few days. Wing cells emerge as intermediaries, facilitating layered structure formation. By day 7, a complete stratified structure forms with all cell types present at the appropriate positions, and by day 15, the tissue achieves a stable homeostatic state with consistent layer organization and thickness.

**Fig 6.**
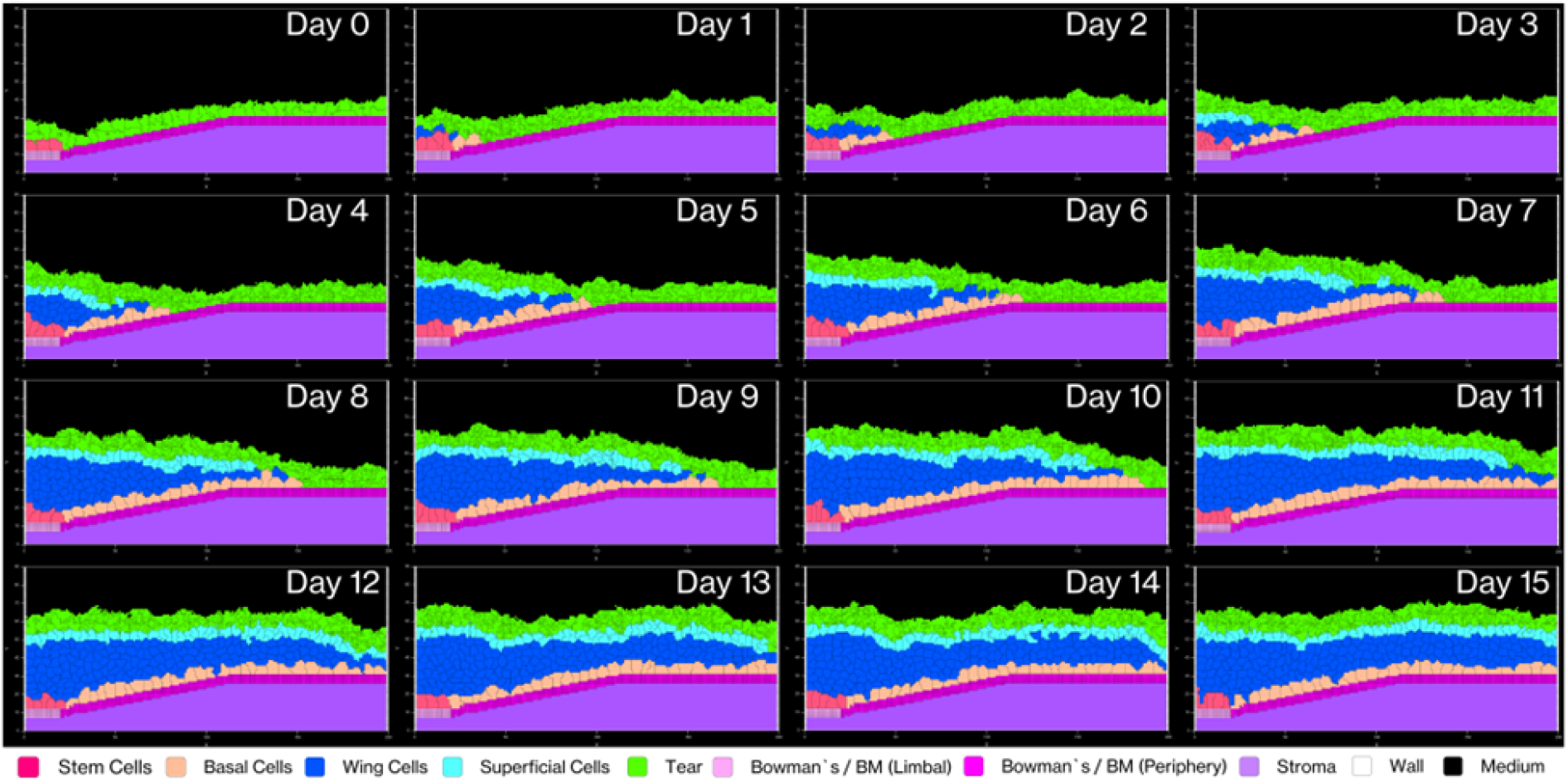
Corneal epithelium development over 15 simulated days. The progression of stratified epithelial layers emerging from an initial population of limbal epithelial stem cells (LESCs, rose-red), the tear film, and the combined Bowman’s layer and EpBM. We begin our simulation with only LESCs at the limbal region (left), followed by centripetal migration and differentiation of basal cells. By day 7, we observe a complete stratified structure forming with all cell types present in the limbus area at nearly the correct height. The final configuration at day 15 demonstrates a stable homeostatic state with consistent layer organization and thickness across the epithelium, characteristic of normal corneal tissue architecture.

This timeline is particularly noteworthy considering the model implements biologically constrained cell cycle times with a minimum doubling interval of 8 hours under optimal conditions. Previous simulations used corneal epithelial cell doubling times ranging from 6 hours to 16 days (29), our model consistently produces tissue-level organization within 7-14 days, aligning with known corneal epithelial renewal periods observed in vivo (84).

### 3.2 Long-term Stability and Tissue Architecture

Our simulation demonstrates long-term stability when run with calibrated parameters (detailed in Supplementary Information Tables S8.1–S8.4). Figure 7 shows the maintenance of consistent cell population dynamics across multiple simulation replicates over six months of simulated time. The frequency distribution of cell counts exhibits a bell-shaped curve, reflecting natural biological variability within controlled parameters. Q-Q analysis confirms that cell count distributions align closely with normal distributions, demonstrating the reproducibility and statistical reliability of our model’s homeostatic state.

**Fig 7.**
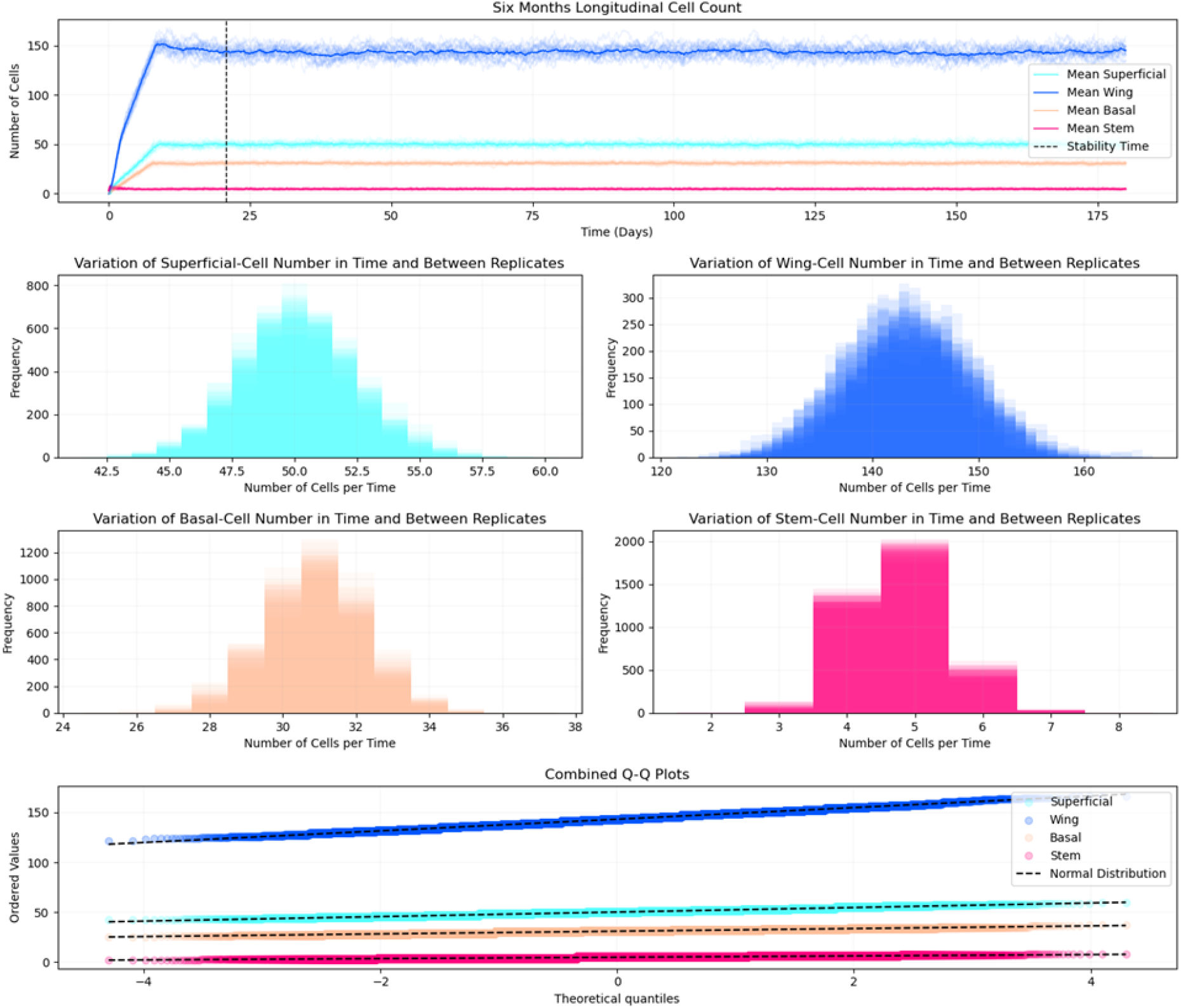
Quantitative analysis of cell population dynamics over six months of simulation. (Top) Mean counts of different cell types (superficial, wing, basal, and stem cells) with our vertical dashed line indicating stability time. (Middle) Display frequency distribution histograms revealing the variation in cell numbers across our replicates for each cell type: superficial cells (cyan), wing cells (blue), basal cells (peach), and stem cells (pink). (Bottom) Presents combined Q-Q plots demonstrating the normality of cell count distributions for all cell types, with dashed lines representing theoretical normal distributions. We observe the close alignment of data points with the normal distribution lines, indicating consistent and stable tissue architecture across our replicates.

We validate our model’s biological accuracy through two complementary approaches for measuring tissue thickness. The Center of Mass Difference method calculates the distance between the deepest cell layer (basal and stem cells) and the superficial cell layer, providing a direct measurement of epithelial thickness. This method shows that the tissue stabilizes at approximately 50 μm, aligning with human corneal epithelial measurements that typically range from 50 to 52 μm (60).

To assess regional variations, we employ Relative Position Tracking using the segmentation approach shown in Figure 8. By dividing the tissue into ten equal segments along the X-axis (0-200 μm) from limbus to periphery, we track the positions of superficial cells within each segment over six months of simulated time. This analysis reveals remarkable consistency across all segments, with thickness variations showing a standard deviation of only 2 μm, indicating highly uniform tissue architecture. Our measurements also demonstrate a characteristic thickness gradient from limbus to periphery, further validating our model’s ability to maintain physiologically relevant tissue architecture.

**Fig 8.**
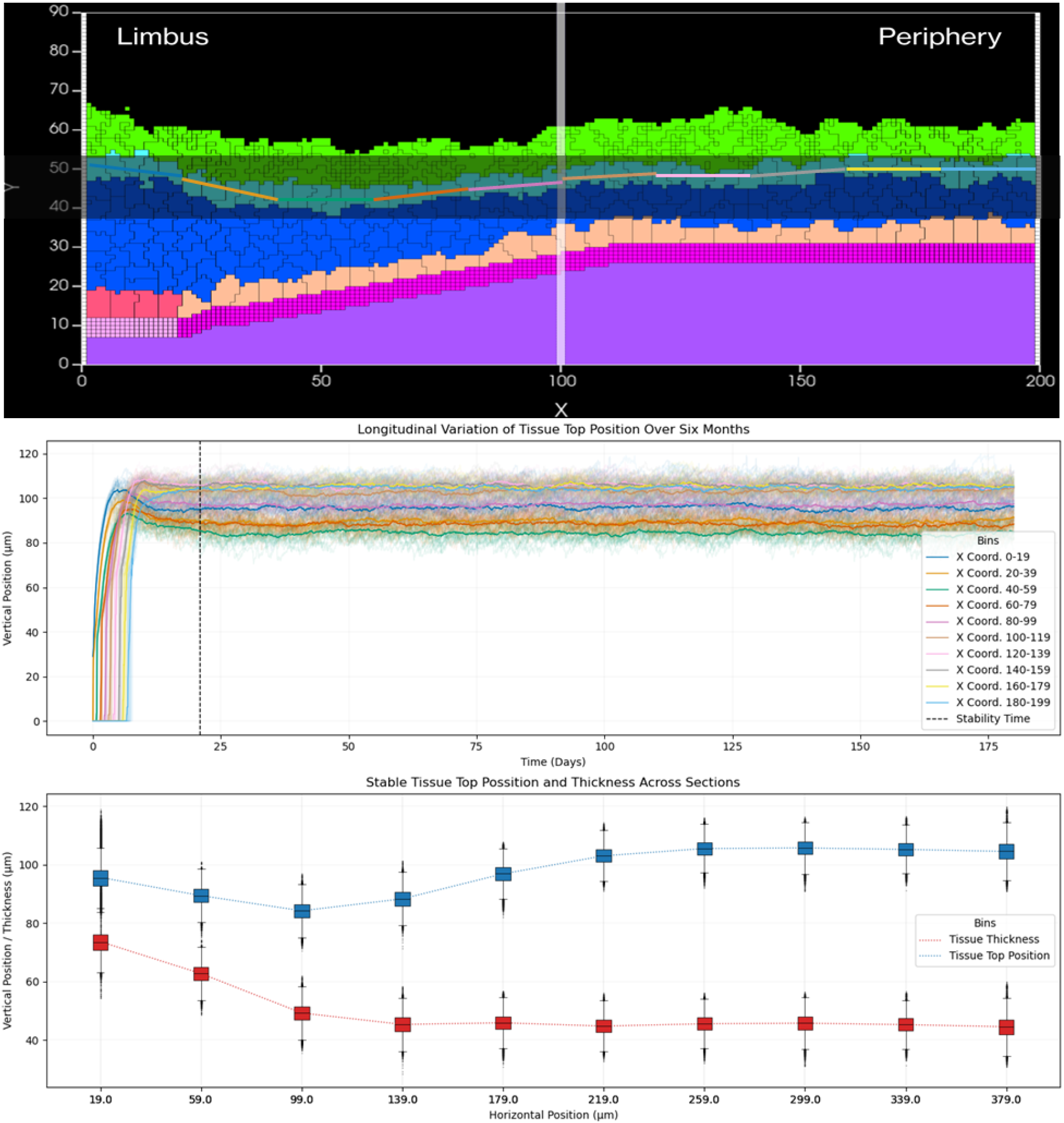
Analysis of tissue thickness and position stability through spatial segmentation. Representative cross-section of the simulated corneal epithelium demonstrating our segmentation strategy for measurements. We divide the tissue into ten equal segments along the X-axis (0-200 μm), from limbus to periphery. Starting from the limbus (left), we create segments corresponding to the X-coordinates shown in the legend of the longitudinal plots: 0-19 (blue), 20-39 (golden-yellow), 40-59 (green), 60-79 (orange), 80-99 (pink), 100-119 (brown), 120-139 (light-pink), 140-159 (grey), 160-179 (yellow-green), and 180-199 (light-blue) in pixels. The plot shows a longitudinal tracking of tissue surface position over six months, where each colored line represents the averaged center of mass position of superficial cells within each corresponding X-coordinate segment. We mark the tissue stability achievement time point with a vertical dashed line. Below we present our quantitative analysis of stable tissue configuration showing both tissue thickness (red squares) and top position measurements (blue squares) with error bars representing temporal variation. Our measurements demonstrate the characteristic thickness gradient from limbus to periphery, validating our model’s ability to maintain physiologically relevant tissue architecture.

### 3.3 Cellular Turnover Dynamics

V-Cornea accurately captures the differential cellular turnover rates observed in biological corneal epithelium. Peripheral corneal regions exhibit faster turnover, with nearly complete cellular substitution occurring approximately every seven days, matching literature reports that indicate corneal epithelium renewal every 7 to 10 days (89–91). In contrast, the limbus region demonstrates a more gradual renewal rate, aligning with a 14-day turnover period.

This differential turnover pattern reflects natural corneal epithelial function, where limbal stem cells serve as a continuous source for epithelial regeneration while maintaining their own population through controlled division rates (7,84). We maintain this realistic dynamic in our model through a balanced equilibrium between cell proliferation and death, even over extended simulation periods. This temporal heterogeneity emerges naturally from our model despite implementing consistent rules across all cells, demonstrating how V-Cornea captures complex tissue behaviors through simple cellular mechanisms.

### 3.4 Injury Simulation and Recovery

To evaluate our model’s capacity to simulate tissue resilience and recovery, we introduced injuries of varying depths based on the classification system established by Scott et al. (2010). Our post-injury simulations revealed distinct recovery patterns that varied with injury depth, as illustrated in Figure 9.

**Fig 9.**
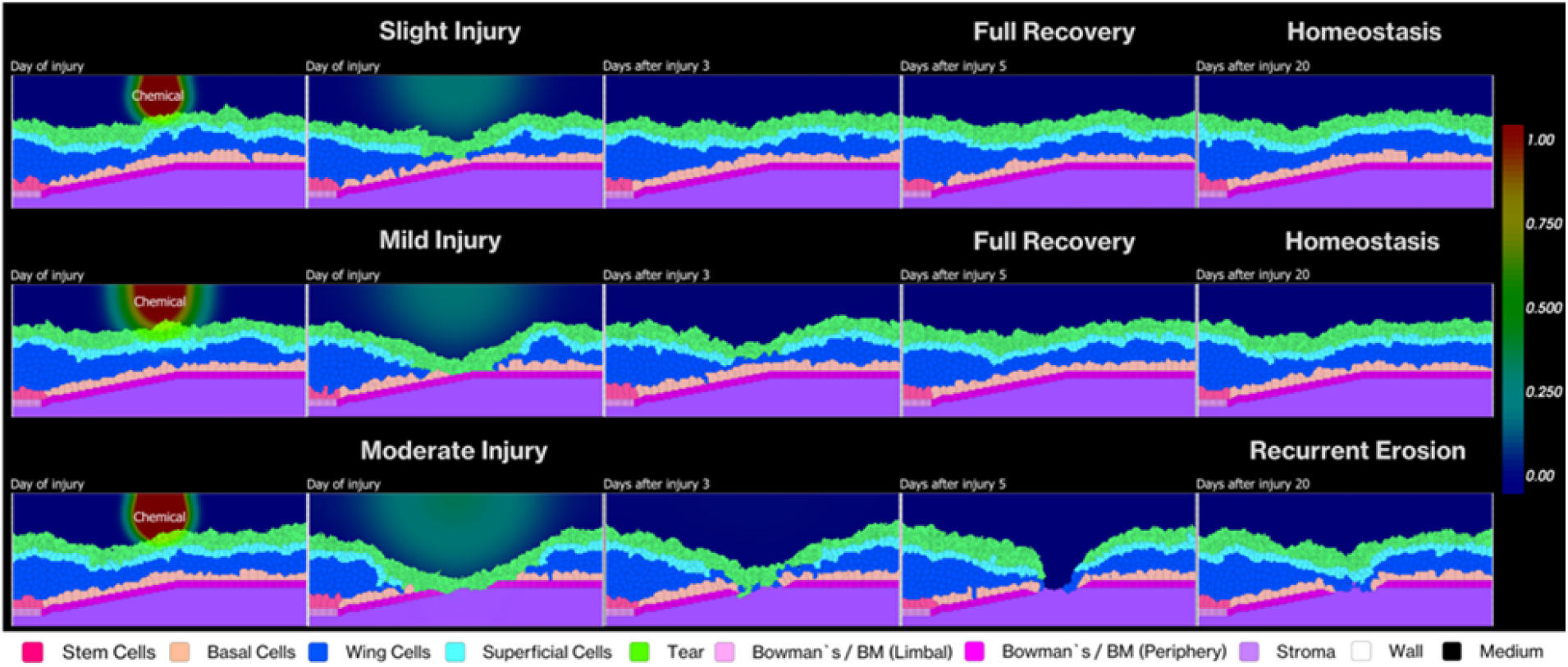
Temporal progression of corneal epithelial recovery following chemical injuries of varying severity. We demonstrate three distinct injury patterns and their recovery over 20 days in our simulation: Slight injury was generated setting a pixel at position (100,75) with concentration 750 a.u. (top row) affecting only superficial and wing cells shows complete recovery and maintained homeostasis; On Mild injury (middle row) was caused by changing the concentration to 1500 a.u. extending to basal cells also achieves full recovery and homeostasis; Increasing the concentration of 2500 a.u. Moderate injury (bottom row) penetrating through to the stroma results in recurrent erosion due to compromised basement membrane. The color gradient bar (right) indicates chemical concentration from 0.00 to 1.00. Each row shows progression from day of injury through days 3, 5 and 20 post-injury.

For slight and mild injuries limited to the epithelial layers, we observed complete recovery within 3-5 days post-injury. The simulation captured the immediate response to cell loss—increased EGF concentration within the epithelium that promoted basal cell proliferation and facilitated rapid wound closure. This timeline aligns with clinical studies showing that superficial corneal abrasions in humans typically resolve within 48-72 hours (7,92).

In contrast, moderate injuries that extended into the stroma and compromised the basement membrane produced markedly different outcomes. These deeper injuries resulted in incomplete recovery with persistent disruptions in the epithelial layer, resembling clinical patterns of recurrent corneal erosions (RCE). Figure 10 quantitatively demonstrates this difference—slight and mild injuries show cell populations returning to pre-injury levels within 5-7 days with stable homeostasis thereafter, while moderate injuries lead to sustained perturbations in wing and superficial cell populations over six months of simulation.

**Fig 10.**
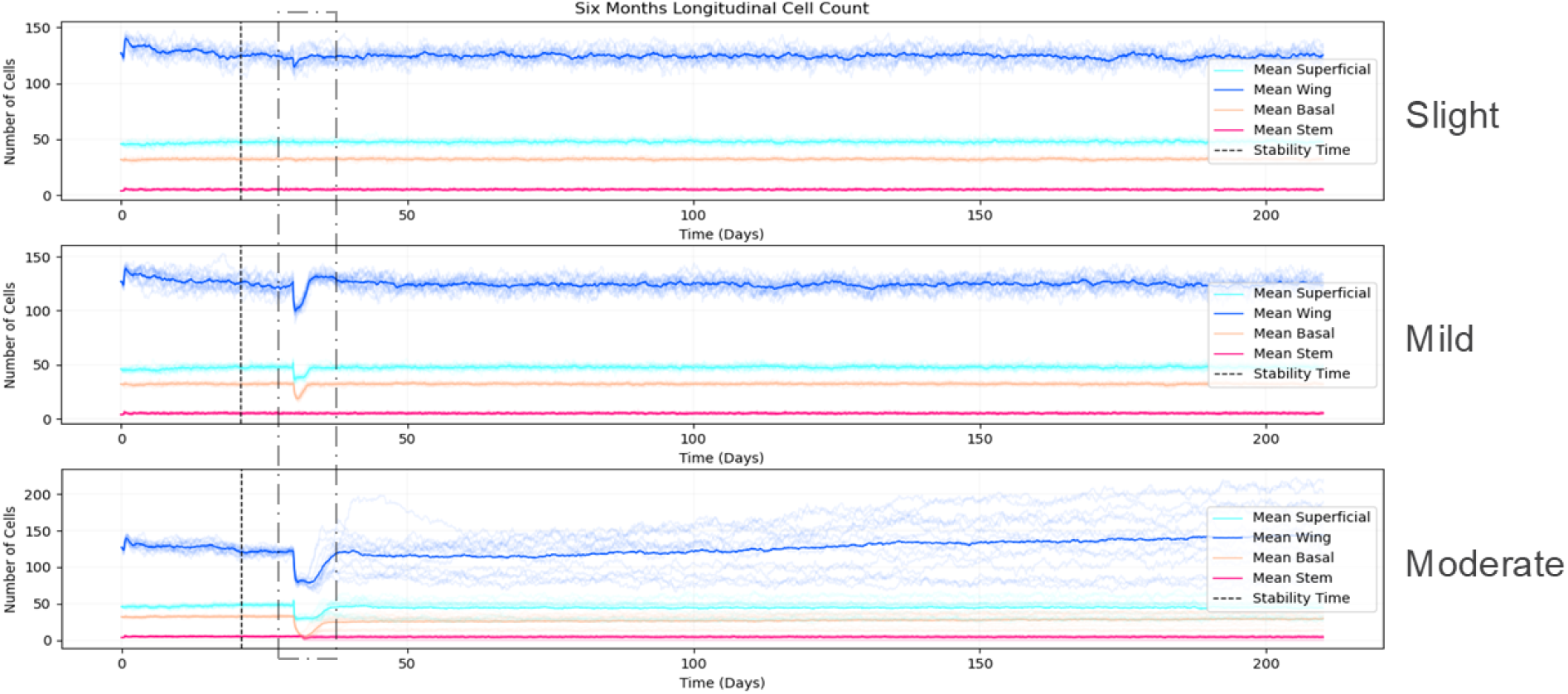
Comparative quantitative analysis of cell population dynamics six months following injuries. Tracking cell counts longitudinally over 6 months after injury, to show tissue response to slight (top), mild (middle), and moderate (bottom) injuries. Mean cell counts for different cell types are the solid lines: superficial cells (cyan), wing cells (blue), basal cells (peach-orange), and stem cells (rose-red). We indicate standard deviation across multiple simulations with shaded areas. We mark the stability time (pre-injury) and injury time points with vertical dashed lines. Following slight and mild injuries, we observe cell populations returning to pre-injury levels within 5-7 days, maintaining stable homeostasis thereafter. In contrast, we find moderate injury leads to sustained perturbation in cell numbers, particularly in wing and superficial cell populations, indicating incomplete recovery. This behavior mirrors clinical observations of recurrent corneal erosions following deep injuries that compromise basement membrane integrity.

This emergent RCE-like behavior stems from a specific mechanism in our model. When the basement membrane becomes disrupted, basal epithelial cells fail to establish stable adhesion to the underlying stroma, resulting in repeated epithelial detachment after initial closure. The effect arises because the basement membrane serves dual critical functions: providing a substrate for epithelial cell adhesion through hemidesmosomes, while simultaneously maintaining basal cells in an undifferentiated state (48). When basal cells lose contact with the basement membrane, they undergo premature terminal differentiation, compromising epithelial integrity.

### 3.5 Model Validation and Relationship to Clinical Conditions

The alignment between our simulation results and empirical data serves as validation of the model’s biological relevance. The recovery period of 3-5 days for superficial injuries parallels findings from multiple clinical studies (7,93), consistent with the natural turnover rate of corneal epithelial cells (84).

More significant is our model’s ability to reproduce key aspects of recurrent corneal erosions, despite not explicitly programming this pathology. In clinical RCE, basement membrane damage disrupts hemidesmosome formation and adhesion complexes, leading to persistent epithelial defects (72,73,94). The molecular basis for this disruption involves processes such as improper processing of laminin-5, a key basement membrane component essential for stable epithelial adhesion (95).

Our current model does not incorporate specific molecular mechanisms of basement membrane regeneration, which would involve coordinated interaction between basal epithelial cells and stromal keratocytes (84). This limitation actually provides an unintended benefit—creating a system where we can study the consequences of impaired basement membrane function in isolation from repair mechanisms.

The emergence of RCE-like features highlights how fundamental the basement membrane is to corneal epithelial stability. While our findings cannot directly inform the pathogenesis of clinical RCE, our model demonstrates that compromised basement membrane integrity alone is sufficient to produce recurrent epithelial breakdown, independent of other factors that may contribute to the clinical condition. This computational finding supplements existing literature on basement membrane dysfunction in corneal pathologies (96), offering a simplified system where the isolated effects of basement membrane compromise can be observed.

Our focus on depth-based injury patterns rather than specific injury mechanisms reflects the clinical observation that healing time correlates strongly with injury depth regardless of the initial insult type. This consistency suggests that our model captures fundamental biological processes governing corneal wound healing. Future iterations could incorporate mechanisms for basement membrane regeneration to better distinguish between model limitations and accurate biological representations, as well as targeted comparisons with chemical-specific clinical data to refine the model’s predictive capabilities.

## 4. Discussion

### 4.1 Interpretation of Findings

V-Cornea demonstrates that complex corneal epithelial behavior can emerge from relatively simple biological rules. Previous modeling approaches have typically addressed isolated aspects of corneal dynamics—mechanical properties through FEA or wound healing through PDEs— whereas our integrated approach illuminates how cellular-level decisions collectively generate sophisticated tissue-level organization. The maintenance of tissue homeostasis over extended simulation periods provides compelling evidence that fundamental corneal epithelial mechanisms can be effectively captured through a limited set of core cellular rules.

Strong alignment between simulated turnover rates and empirical observations (84,89–91,97) validates that these basic rules accurately reflect underlying biological processes. Particularly significant is the successful replication of the mechanistic relationship between basement membrane contact and cell differentiation (45,48), a key regulatory pathway previously uncaptured in computational models. This reproduction of both structural and functional aspects of corneal maintenance demonstrates that essential features of epithelial self-organization can be captured in a simplified rule set.

### 4.2 Response to Injury and Recovery Mechanisms

The V-Cornea model provides valuable insights into mechanistic relationships between tissue damage and recovery by simulating differential healing responses based on injury severity. The successful replication of recovery times for slight and mild injuries validates our implementation of EGF-mediated healing mechanisms (4,5). Particularly noteworthy is the emergent relationship between epithelial barrier disruption and accelerated healing through increased EGF diffusion— a phenomenon that arises naturally from basic cellular rules rather than explicit programming.

More significant, however, is the model’s emergent behavior when simulating basement membrane disruption. What initially appeared as a limitation—the inability to fully heal moderate injuries—actually reveals valuable insights into pathological conditions such as recurrent corneal erosions (72,98,99). These simulations demonstrate that V-Cornea can reproduce clinically relevant outcomes from relatively simple rule sets, making it a valuable platform for both theoretical and applied research. The model provides a framework that experimentalists and clinicians can use to investigate mechanisms, test hypotheses, and potentially develop therapeutic interventions for corneal pathologies without exclusive reliance on animal models or extensive clinical trials.

These findings highlight a crucial direction for future model development: incorporating the dynamic interplay between epithelial cells and keratocytes in basement membrane regeneration. Such enhancements would improve predictive capabilities for moderate injuries while potentially offering new insights into therapeutic approaches for recurrent corneal erosions. The emergent behaviors observed in V-Cornea demonstrate its potential as a versatile and extensible platform that bridges basic science and clinical applications in corneal research.

V-Cornea serves as a valuable bridge between experimental endpoint measurements and the dynamic progression of healing responses. Its potential applications extend to toxicology, where it could provide a computational framework for eye irritation testing by correlating patterns of initial cellular damage with distinct adverse outcomes. This approach could eventually complement existing testing methods, though significant validation work remains necessary before such applications can meaningfully contribute to reducing animal testing or informing regulatory decisions.

### 4.3 Limitations and Future Directions

Although V-Cornea effectively simulates acute corneal injury response, several biological mechanisms require implementation to enhance its relevance for accidental eye exposure scenarios. Our treatment of the basement membrane as a non-regenerative entity represents a key limitation in accurately predicting recovery from chemical exposures. After acute injury, epithelial cells and stromal keratocytes regenerate the basement membrane through coordinated action (72,95) - a process that, once implemented, would significantly improve predictions for injuries compromising basement membrane integrity.

The current model’s simplified representation of chemical injury mechanisms presents a second limitation. Chemical exposures are currently modeled using generic diffusion patterns and concentration-dependent cell death thresholds, without incorporating chemical-specific modes of action or cellular selectivity. While this approach successfully distinguishes between injury classifications based on depth, it fails to capture the diversity of chemical-tissue interactions observed in real exposures. Different chemicals (surfactants, acids, alkalis, organic solvents) interact with corneal tissue through distinct mechanisms - disrupting cell membranes, denaturing proteins, or altering cellular metabolism. Future iterations would benefit from experimental data on specific chemical reactivity, potency, and cellular targets to better simulate how different chemical classes affect corneal tissue, particularly for consumer products containing multiple ingredients with varying modes of action.

A third critical limitation lies in the simplified representation of tear film dynamics. The current model assumes constant tear production and EGF levels, whereas recent research shows these factors change rapidly following acute chemical exposure (33). Since tear composition significantly influences both initial chemical exposure and subsequent healing response, future versions should incorporate dynamic tear film properties through variable diffusion rates and concentration-dependent cellular responses based on exposure conditions.

The exclusion of stromal and endothelial layers also limits comprehensive assessment of chemical injury responses, particularly for exposures penetrating beyond the epithelium. Without these deeper layers, important acute responses such as keratocyte activation and myofibroblast transformation (7) remain uncaptured. Expanding the model to include these components would enable more comprehensive simulation of tissue responses to chemical exposures of varying severity.

Emerging experimental technologies offer opportunities to significantly advance V-Cornea’s biological relevance. Spatial single-cell RNA-sequencing data collected at different timepoints during acute injury and recovery could reveal critical temporal dynamics of cellular responses and identify key molecular players in wound healing. Such data would enable more sophisticated modeling of cell-state transitions and intercellular signaling networks, particularly for chemical exposures, providing deeper insights into adverse outcome pathways.

Future development priorities for V-Cornea should focus on implementing basement membrane regeneration mechanisms, developing dynamic models of tear film composition, integrating stromal and endothelial tissue layers, and incorporating mechanisms for immediate inflammatory responses to chemical exposure. The modular CompuCell3D implementation provides a flexible framework that researchers can adapt for diverse corneal studies beyond acute chemical exposure. Collaboration with research groups across toxicology, ophthalmology, and basic corneal biology will strengthen V-Cornea’s utility while advancing our broader understanding of corneal biology and pathology.

## 5. Conclusion

V-Cornea effectively simulates complex corneal epithelial behavior using an agent-based modeling approach with biologically-inspired rules. The model successfully reproduces key aspects of corneal epithelial dynamics, including tissue self-organization, maintenance of stable architecture, and physiologically relevant cell turnover rates. Validation against empirical measurements (84,89–91,97), establishes V-Cornea’s ability to capture fundamental homeostatic and recovery processes in corneal epithelium.

The model’s ability to predict differential healing responses based on injury depth represents a particularly significant achievement. V-Cornea demonstrates potential for investigating pathological conditions (72,98,99) through accurate reproduction of healing times for slight and mild injuries (4,5,7), coupled with emergent behavior mimicking recurrent corneal erosions when basement membrane disruption occurs. These capabilities emerge naturally from basic cellular rules, indicating that the model captures essential features of corneal wound healing dynamics.

Through its modular CompuCell3D implementation, V-Cornea provides an extensible framework that researchers can adapt for diverse applications in corneal biology. While the current version does not yet include basement membrane regeneration mechanisms and deeper tissue responses, adding these components represents a priority for future development. We invite collaboration with research groups interested in applying or extending the model’s capabilities for studying basic corneal biology, evaluating chemical toxicant effects, or exploring other aspects of corneal pathology. This open, collaborative approach aims to advance understanding of corneal biology while supporting the development of animal-free methods for assessing corneal responses to injury.

## Declaration of generative AI and AI-assisted technologies in the writing process

During the preparation of this work the author(s) used OpenAI | ChatGPT o1 and Anthropic | Claude 3.7 Sonnet in order to improve the readability and avoid repetition of information. After using this tool/service, the author(s) reviewed and edited the content as needed and take(s) full responsibility for the content of the published article.

## CRediT authorship contribution statement

Joel Vanin: Conceptualization, Methodology, Software, Validation, Formal analysis, Investigation, Data Curation, Writing - Original Draft, Writing - Review & Editing, Visualization

Michael Getz: Methodology, Investigation, Data Curation, Review & Editing

Catherine Mahony: Conceptualization, Investigation, Review & Editing, Supervision, Project administration, Resources, Funding acquisition

Thomas B. Knudsen: Review & Editing, Supervision

James A. Glazier: Conceptualization, Project administration, Review & Editing, Supervision

## Data and Code Availability

All model code, simulation files, scripts for plots, and data used to generate the results presented in this study are openly available in a public repository. The V-Cornea model, implemented in CompuCell3D, along with the scripts for the plots and the data files, can be accessed on GitHub at https://github.com/VaninJoel/vCornea. Additionally, a permanent, versioned archive of the code is also available on Zenodo at https://doi.org/10.5281/zenodo.16764319. All other data supporting the findings of this study are available within the article and its Supplementary Information files.

## S1. Growth Dynamics Mathematical Formulation

### S1.1 EGF-Dependent Growth (Hill Function)

For basal and stem cell types, the growth factor due to EGF concentration is defined as:

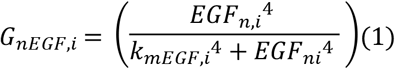

where *EGF*_*ni*_ is the average EGF concentration in cell *i* at time *n*. We define:

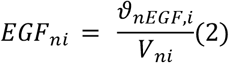

Where

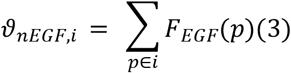

i.e.,*ϑ*_*nEGF,i*_ is the total EGF amount inside cell *i*, obtained by summing the EGF field *F*_*EGF*_ over all lattice pixels p belonging to cell *i. V*_*ni*_ is the cell’s volume (in number of pixels or voxels) at time *n*. The parameter *k*_*mEGF,i*_ is the half-maximal EGF concentration for cell *i*.

### S1.2 Density-Dependent Growth Inhibition (Hill Function)

Cells experience density-dependent growth inhibition described by another Hill-type function:

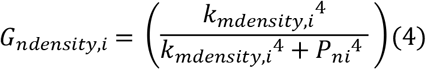

where *P*_*ni*_ is the effective “pressure” inside cell *i* at time *n*, derived from its volume deviation (see below). The parameter *k*_*mdensity,i*_ is the half-maximal pressure for growth inhibition.

### S1.3 Effective Pressure Calculation

The volume energy term in the Hamiltonian is:

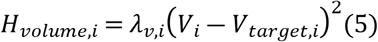

where ***λ***_***v***,***i***_ is the volume constraint parameter for cell *i*; ***V***_***i***_ is the actual (current) volume of the cell ***i***, and ***V***_*target*,***i***_ is the target volume of the cell ***i***.

The effective pressure inside cell *i* is then:

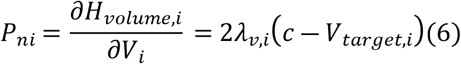

this pressure term reflects how far the cell’s volume is from its target.

### S1.4 Total Growth Rate

Combining the EGF-dependent growth and density-dependent inhibition terms, each cell’s net growth rate is:

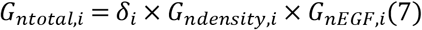

where *δ*_*i*_ is the cell’s intrinsic maximal doubling rate (e.g., *δ*_*basal*_ or *δ*_*stem*_) given in hours. Finally, cell *i* grows by updating its target volume:

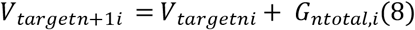

### S1.5 Parameter Fitting for Hill Function Coefficients

The half-maximal concentration parameters (*k*_*mEGF*_ and *k*_*mdensity*_) are critical control points in our growth regulation model, directly influencing proliferation rates in response to both EGF signaling and contact inhibition. To determine appropriate values for these parameters, we employed a multi-objective optimization approach constrained by three key biological criteria:

1. **Epithelial Turnover and Recovery Timeline**: Values were calibrated to ensure that complete epithelium turnover time of 7-14 days, and the recovery of 3-5 days after injury, matched the empirically observed times in healthy corneal epithelium (7,84).
2. **Layer-Specific Cell Density**: Parameter values were adjusted to maintain physiologically relevant cell numbers, 5–7 layers of cells (100).
3. **Epithelial Thickness Maintenance**: We constrained our parameters to maintain stable epithelial thickness of 54 ± 5 μm over extended simulations, consistent with in vivo measurements (100).

We performed an iterative parameter sweep, systematically varying *k*_*mEGF*_ from 1.0 to 10.0 and *k*_*mdensity*_) from 10.0 to 200.0 across multiple simulation runs. The optimal values (*k*_*mEGF, stem*_ = 3.5, *k*_*mEGF, basal*_ = 7.0 and *k*_*mdensity*_ = 125.0 for both) produced the closest match to all three biological constraints simultaneously.

### S1.6 Mitosis Rule

Mitosis in our model is governed by a volume-threshold mechanism that initiates cell division once a cell accumulates sufficient biomass.

Formally, for any proliferative cell *i* (stem or basal), division occurs when:

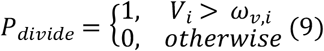

where *V*_*i*_ represents the current volume of cell, and*ω*_*v,i*_ is the cell type-specific volume threshold for division. We set 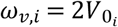 where 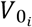 is the initial cell volume (25 pixels or approximately 100 μm^2^).

When a cell reaches this threshold, we implement division through the following algorithm:

1. Identify the division plane orientation: For limbal epithelial stem cells (LESCs), we orient the division plane to direct daughter cells centripetally, following observed patterns that maintain stem cell populations while producing committed progenitors (64). For basal cells, we select a random division plane to promote even tissue distribution (48).
2. Create two daughter cells with equal volumes:

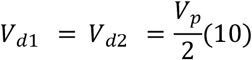

where *V*_*d*1_ and *V*_*d*2_ are the volumes of the daughter cells, and *V*_*p*_ is the parent cell’s volume at division.
3. Reset target volumes for daughter cells to their initial type-specific values:

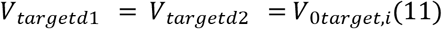
4. Preserve cell type-specific properties: Each LESC division produces one stem cell and one basal cell, maintaining the stem cell pool while generating committed progenitors (83).

Basal cell divisions produce two identical basal cells, both capable of further proliferation.

## S2. Differentiation Rules Mathematical Formulation

### S2.1 Stem to Basal Differentiation

A limbal epithelial stem cells (LESCs) differentiates into basal cells it loses contact with limbal Bowman’s layer /Epithelial Basement Membrane (EpBM):

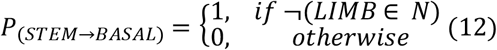

here *N* is the set of neighboring cell types, ¬*LIMB* ∈ *N* means the LIMB type is not in the neighbor set.

### S2.2 Basal to Wing Differentiation

Basal cells differentiate to wing cells based on their contact area with Bowman’s layer/Epithelial Basement Membrane (EpBM):

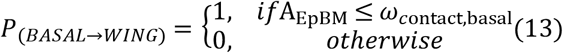

### S2.3 Wing to Superficial Differentiation

A wing cell differentiates into a superficial cell once it contacts the surface (tear film) and loses contact with deeper layers. The transition is:

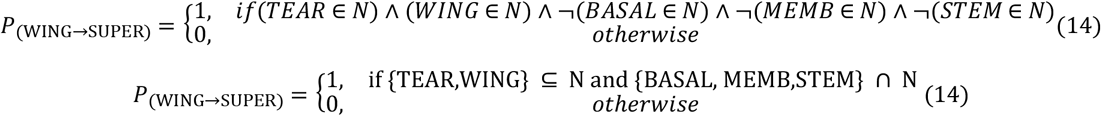

All these transitions are deterministic (P=1 or P=0) and occur immediately once the condition is met.

## S3. Movement Implementation in the Cellular Potts Model

### S3.1 Hamiltonian Definition

Cell movement is governed by the minimization of the system’s Hamiltonian:

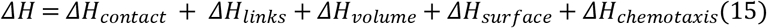

### S3.2 Contact and Links Energy Terms

The contact energy is calculated as:

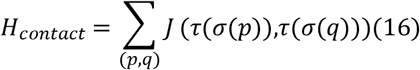

where *J*(*τ*_1_,*τ*_2_) is the contact energy between cell types τ_1_ and τ_2_ and *σ*(*p*) is the cell occupying lattice site *p. τ*(*σ*) gives the cell type of cell σ.

Each link is modeled as a Hooke-type (spring) interaction between the centers of mass of two cells (or between one cell and a “wall” cell). These springs can be created, updated (e.g., to maintain a certain tension), or removed depending on local conditions in the simulation.

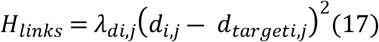

Where *λ*_*di,j*_is the link`s spring constant, *d*_*i,j*_ is the current distance (in voxels) between the cell centers, *d*_*targeti,j*_is the links equilibrium length.

From this energy, the tension (force) on each cell due to the link is:

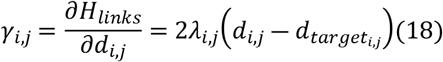

We enforce a constant tension on the superficial cells layer by automatically adjusting either *λ*_*i,j*_ or 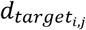.

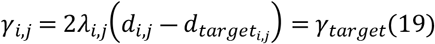

#### Case of Adjusting *λ*_*i,j*_

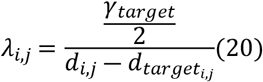

#### Case of Adjusting 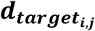

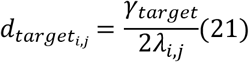

Depending on the user’s “AutoAdjustLinks” and “Lambda_link_adjustment” settings, the code picks one of these two strategies to keep *γ*_*i,j*_ at the desired tension.

### S3.3 Volume and Surface Constraints

Volume and surface energy terms are:

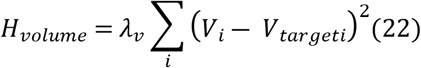

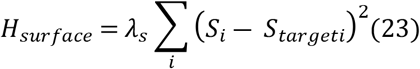

Where *λ*_*v*_, *λ*_*s*_ are volume/surface coefficients; *V*_*i*_, *S*_*i*_ are the current volume/surface area. 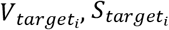 are the respective targets.

### S3.4 Chemotaxis Term

The chemotactic energy contribution is:

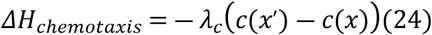

Where *λ*_*c*_ is the chemotactic coefficient, and *c*(*x*) is the chemical concentration at position *x*. A proposed copy from *x* to *x*^′^ changes the Hamiltonian according to *ΔH*.

### S3.5 Movement Algorithm

1. Randomly choose a lattice pixel *x* and one of its neighbors *x*^′^.
2. Compute ΔH for copying σ(x)→σ(x′).
3. Accept with probability

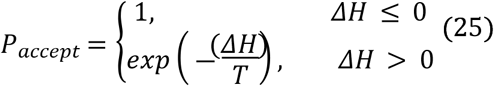

where T is the simulation temperature parameter.

## S4. Cell Death and Sloughing Mathematical Formulation

### S4.1 Natural Cell Death (Sloughing)

For superficial cells in contact with tear film, the probability of sloughing is:

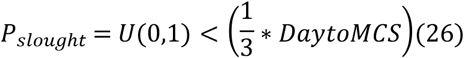

where *U*(0,1) is a uniform random sample in [0,1], and *DaytoMCS* is the factor converting between days and Monte Carlo Steps. The multiplier 1/3 is calibrated so that the average layer transit time is about 1.75 days out of a total 7–14 day turnover cycle.

### S4.2 Volume-Based Death Condition

For superficial cells:

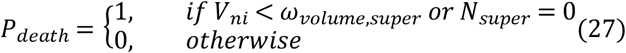

where *V*_*ncell,i*_ is current cell volume, *ω*_*volume,super*_ = 15 (minimum viable cell size), *N*_*super*_ is the number of neighboring superficial cells.

### S4.3 Volume Reduction During Death

Dying cells shrink to zero volume:

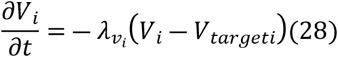

where for a dying cell, 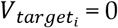 and 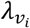 is increased to 1000 for rapid shrinkage.

### S4.4 Injury Implementation

#### A. Ablation Injuries

Ablation injuries are simulated by the instantaneous removal of cells within a defined circular region. Any part of the cell included in the circular area will be selected to be part of the cells set to be removed, defined as:

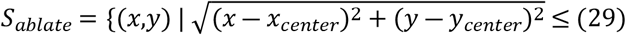

All cells intersecting *S*_*ablate*_ are converted to “tear” type:

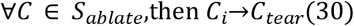

simulating the removal of epithelium and its replacement by tear fluid in the wound, as observed in corneal wound healing (Wilson et al., 1999).

#### B. Chemical Injuries

Chemical injuries are modeled through a reaction-diffusion equation:

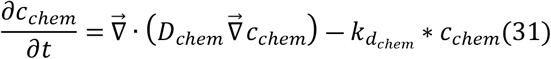

where *c*_*c*h*em*_(*x,y,t*) is chemical concentration, *D*_*c*h*em*_ is the diffusion coefficient that is dependent on localization, 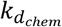 is degradation rate.

With two initial-condition types:

**Gaussian Pulse** (droplet or localized exposure):

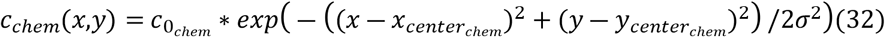

where 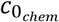 is initial peak concentration, *σ* is distribution width parameter, 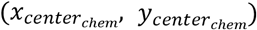 is exposure center point.

**Uniform Distribution** (coating or widespread exposure):

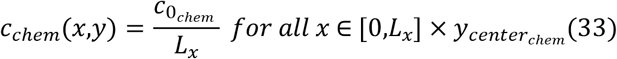

where *L*_*x*_ are the whole lattice dimension in 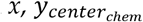 is the given height, and 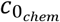 is initial concentration.

### S4.5 Cell Death Conditions

Cells die if their mean chemical concentration exceeds a threshold. Let

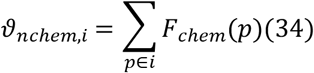

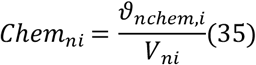

*F*_*c*h*em*_(*p*) is the chemical field value at pixel *p* within the cell, *ϑ*_*nc*h*em,i*_ is the total amount of chemical observed by the cell *i* at time *n, V*_*ni*_ is the volume of the cell *i*, lastly *C*h*em*_*ni*_ is the average concentration of chemical at cell *i* at time *n*. Which define the set of doomed cells in:

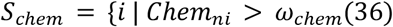

All cells in *S*_*c*h*em*_, undergo rapid volume collapse:

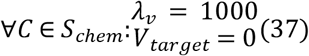

## S5. EGF Dynamics Mathematical Formulation

### S5.1 Spatiotemporal Evolution

The EGF concentration field *c*_*EGF*_(*x,y,t*) evolves according to the reaction-diffusion equation:

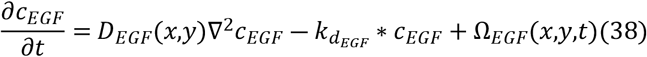

where *D*_*EGF*_(*x,y*) is the cell-type dependent diffusion coefficient, 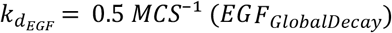 is the global decay rate, Ω_*EGF*_(*x,y,t*) represents the net source/sink terms.

### S5.2 Cell-Type Dependent Transport

EGF diffusion coefficient varies by cell type:

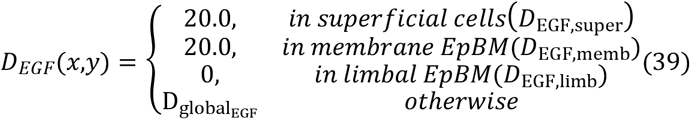

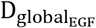 global diffusion constant for EGF was set at 186 voxels^2^/MCS based on experimental measurements adapted to our simulation parameters. We initially calculated the diffusion coefficient from rat brain tissue studies (101), yielding 466.2 voxels^2^/MCS after converting to our spatial (1 voxel = 2 μm) and temporal (1 hour = 10 MCS) scales. However, this value would cause unrealistically rapid equilibration across our simulation domain (200 × 90 voxels).

We therefore considered alternative data from GelMA hydrogel experiments (87), which reported a more restricted EGF diffusion coefficient (2.5 × 10^−8^ cm^2^/s), equivalent to 225 voxels^2^/MCS in our units. Our selected value of 186 voxels^2^/MCS closely approximates this experimentally determined rate for EGF in dense extracellular environments while remaining computationally feasible. This parameter allows for biologically relevant gradient formation, providing an appropriate balance between physiological accuracy and computational efficiency.

### S5.3 Source and Sink Terms

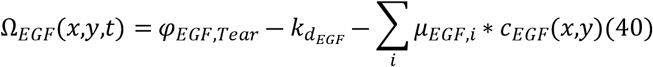

where *φ*_*EGF,tear*_ = 1.0 is the constant secretion rate by tear cells. And *μ*_*EGF,i*_ are cell-type specific uptake rates *μ*_*EGF,basal*_ = 0.0 for basal cells, *μ*_*EGF,stem*_ = 0.0 for stem cells, *μ*_*EGF,super*_ = 0.0 for superficial cells, *μ*_*EGF,wing*_ = 0.0 for wing cells, since we are using a increase global decay as a surrogate to these individual cells uptakes for simplicity.

The global decay constant for EGF 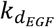 was set to 0.5 per MCS to account for multiple biological processes affecting EGF availability. Using published data on EGF circulating half-life ranging from 42 to 114 minutes (88), we calculated corresponding decay constants between 0.099 and 0.036 per MCS in our simulation units (1 hour = 10 MCS).

Our implemented decay value (0.5) is intentionally higher than these physiological baseline values to implicitly incorporate additional processes not explicitly modeled, including cellular uptake, receptor-mediated endocytosis, proteolytic degradation, and sequestration by extracellular matrix components. This higher decay rate also ensures appropriate diffusion length scales within our simulation domain.

### S5.4 Boundary Conditions

#### Physical Domain Structure

The simulation domain is bounded by non-diffusive wall cells at

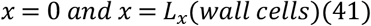

where wall cells have *D*_*EGF,wall*_ = 0 (no diffusion). These cells create effective no-flux boundaries by blocking EGF transport.

#### Mathematical Boundary Conditions

##### Horizontal Boundaries

Effective no-flux condition due to combination of:

1. Zero-derivative boundary condition at domain edges:

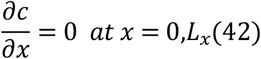
2. Impermeable wall cells creating physical barriers:

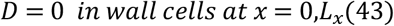

##### Vertical Boundaries

Fixed concentration at top and bottom:

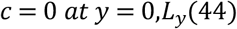

This configuration creates a physically confined system where:

EGF cannot escape through the lateral boundaries (wall cells + no-flux conditions) Top and bottom boundaries maintain zero concentration

Effective transport is confined to the region between wall cells

### S5.5 Spatial and Temporal Scales

Spatial:

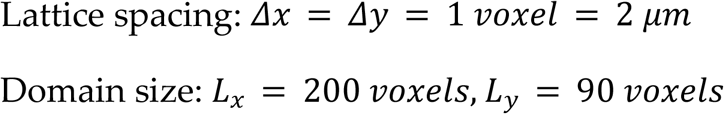

Temporal:

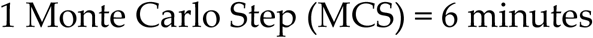

**Table S1.** Stem cells behavior signal relationship.

| Agent Type | Behavior | Form | $\frac{Min}{Max}$ | Signal(s) | Effect(s) | Params |
| --- | --- | --- | --- | --- | --- | --- |
| Stem | Growth (Eq. 8) | Multiplicative Hill (Eq. 7) | $\frac{0}{\delta_{stem}}$ | EGF (Eq. 1) | Increase | Half max: $k_{mEGF,stem}$<br>Hill power: 4 |
| | | | | Pressure (Eq. 4) | Decrease | Half max: $k_{mdensity,stem}$<br>Hill power: 4 |
| | Differentiation to Basal (Eq. 12) | Boolean Conditional | 0/1 | Contact with Limbal BM | Disallow | $\omega_{contact,stem} = 1$ voxel |
| | Mitosis | Boolean Conditional | 0/1 | Cell Volume | Allow | $\omega_{v,stem} = 2V_{0target,stem}$ |
| | Movement (Boltzmann Acceptance Eq. 25) | Contact Energy (Eq. 16) | $\frac{5}{20}$ | Cell Neighbor | Energy Contribution | Table S6 energies |
| | | Volume (Eq. 22) | $\frac{-\infty}{+\infty}$ | Cell Volume | Energy Contribution | $\lambda_{0v,stem}=2.0,$<br>$V_{0target,stem}=25.0$ |
| | | Surface Area (Eq. 23) | $\frac{-\infty}{+\infty}$ | Cell Surface | Energy Contribution | $\lambda_{0s,stem}=2.0,$<br>$S_{0target,stem}=18.0$ |
| | | Chemotaxis (Eq. 24) | $\frac{-\infty}{+\infty}$ | Concentration Gradient | Increase Energy Contribution | $\lambda_{0chemoMbias,stem} = 100$ |
| | Apoptosis | Boolean Conditional (Eq. 36) | 0/1 | Chemical Concentration | Allow | $\omega_{chem}$ |

**Table S2.**
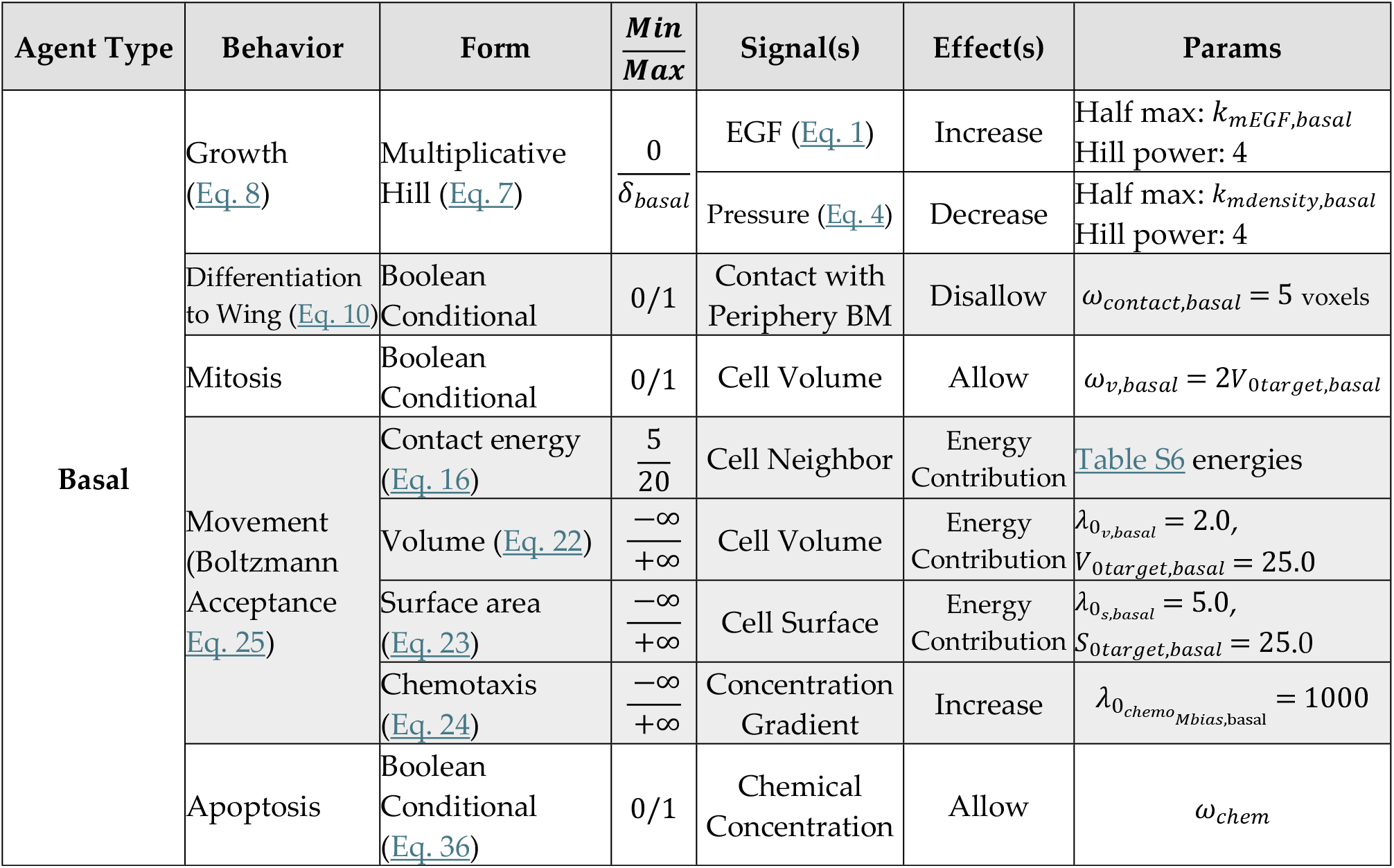
Basal cells behavior signal relationship.

**Table S3.**
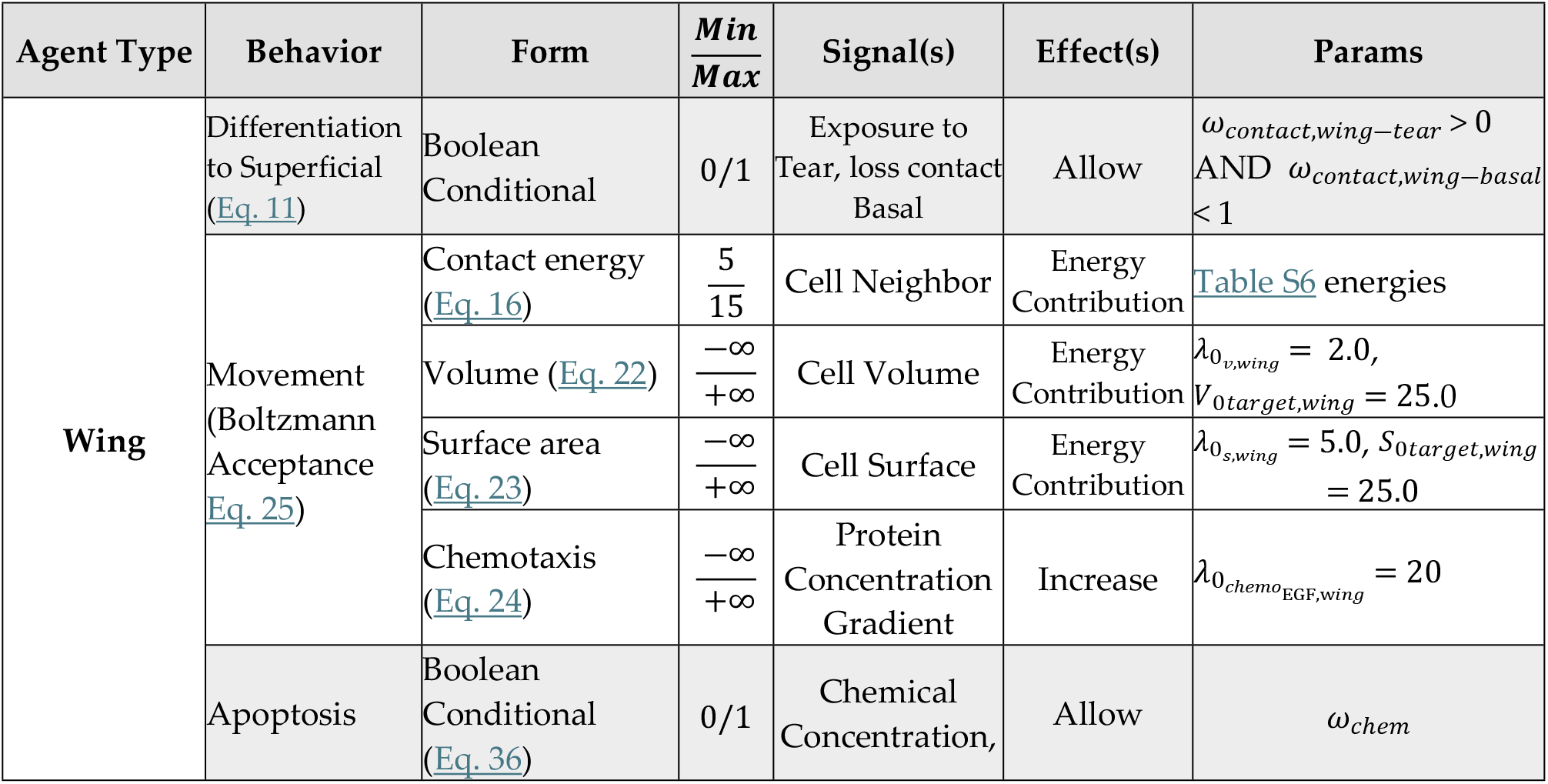
Wing cells behavior signal relationship.

**Table S4.** Superficial cells behavior signal relationship.

| Agent Type | Behavior | Form | $\frac{Min}{Max}$ | Signal(s) | Effect(s) | Params |
| --- | --- | --- | --- | --- | --- | --- |
| Superficial | Movement (Boltzmann Acceptance Eq. 25) | Contact energy (Eq. 16) | $\frac{2}{15}$ | Cell Neighbor | Energy Contribution | Table S6 energies |
| | | Volume (Eq. 22) | $\frac{-\infty}{+\infty}$ | Cell Volume | Energy Contribution | $\lambda_{0,v,super} = 2.0$, $V_{0target,super} = 25.0$ |
| | | Surface area (Eq. 23) | $\frac{-\infty}{+\infty}$ | Cell Surface | Energy Contribution | $\lambda_{0,s,super} = 5.0$, $S_{0target,super} = 25.0$ |
| | | Links (Eq. 17) | $\frac{-\infty}{+\infty}$ | Center of Mass Distance | Energy Contribution | $\lambda_{0link,super} = 50$<br>$L_{0targetlink,super} = 3$<br>$L_{mlink,super} = 1000$<br>$L_{0targetlink,wall} = 3$<br>$\lambda_{0link,wall} = 50$<br>$L_{mlink,wall} = 1000$ |
| | Apoptosis | Boolean Conditional (Eq. 36) | 0/1 | Chemical Concentration | Allow | $\omega_{chem}$ |
| | | Sloughing (Eq. 23) | 0/1 | Probability Draw | Allow | $\left(\frac{1}{3} * DaytoMCS\right)$ |

**Table S5.** Tear and Bowman`s/EpBM behavior signal relationship.

| Agent Type | Behavior | Form | $\frac{Min}{Max}$ | Signal(s) | Effect(s) | Params |
| --- | --- | --- | --- | --- | --- | --- |
| Tear | Movement (Boltzmann Acceptance Eq. 25) | Contact energy (Eq. 16) | $\frac{0.1}{10}$ | Cell Neighbor | Energy Contribution | Table S6 energies |
| | | Volume (Eq. 22) | $\frac{-\infty}{+\infty}$ | Cell Volume | Energy Contribution | $\lambda_{0,v,tear}=1.0, V_{0target,tear}=50.0$ |
| | Secrete EGF | Constant | 1 | — | — | $\varphi_{EGFtear}$ |
| Bowman`s/EpBM | Destructible | Boolean Conditional (Eq. 36) | 0/1 | Chemical Concentration | Allow | $\omega_{chem}$ |

**Table S6.** Contact energy values (arbitrary units)

| Cell Type | Medium | STEM | LIMB | BASAL | WING | SUPER | MEMB | STROMA | WALL | TEAR |
| --- | --- | --- | --- | --- | --- | --- | --- | --- | --- | --- |
| Medium | 10.0 | 10.0 | 10.0 | 10.0 | 10.0 | 10.0 | 10.0 | 10.0 | 5.0 | 5.0 |
| STEM |  | 10.0 | 5.0 | 10.0 | 10.0 | 10.0 | 5.0 | 20.0 | 10.0 | 5.0 |
| LIMB |  |  | 10.0 | 10.0 | 10.0 | 10.0 | 10.0 | 10.0 | 10.0 | 5.0 |
| BASAL |  |  |  | 10.0 | 10.0 | 15.0 | 5.0 | 20.0 | 10.0 | 5.0 |
| WING |  |  |  |  | 5.0 | 6.0 | 15.0 | 10.0 | 10.0 | 5.0 |
| SUPER |  |  |  |  |  | 2.0 | 15.0 | 10.0 | 5.0 | 2.0 |
| MEMB |  |  |  |  |  |  | 10.0 | 10.0 | 10.0 | 2.0 |
| STROMA |  |  |  |  |  |  |  | 10.0 | 10.0 | 10.0 |
| WALL |  |  |  |  |  |  |  |  | 10.0 | 5.0 |
| TEAR |  |  |  |  |  |  |  |  |  | 0.1 |

**Table S7.**
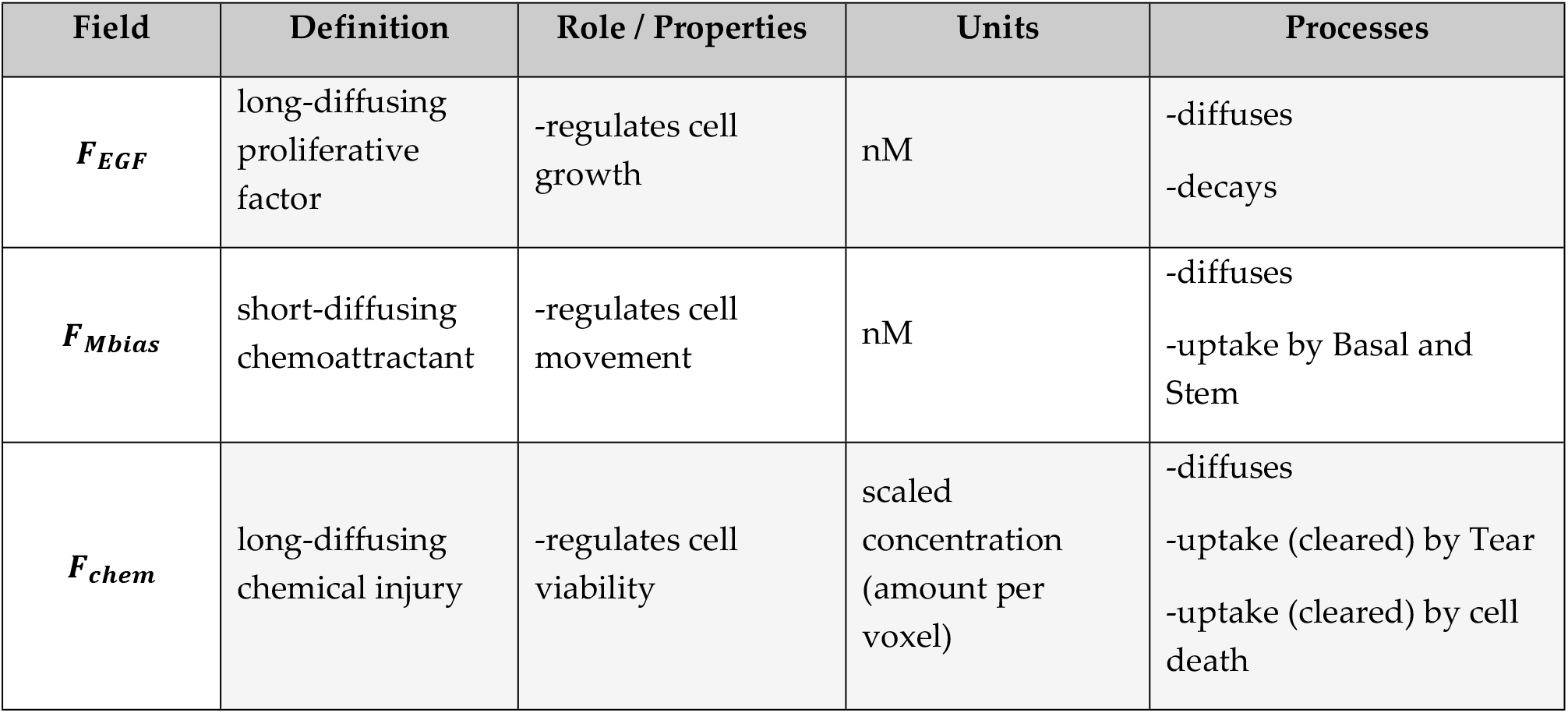
Fields descriptions.

**Table S8.1.** Initial parameters (Cell Growth and Mechanical Constraints)

| Parameter | Symbol | Simulation Value | Literature Ref. | Description |
| --- | --- | --- | --- | --- |
| **Stem Cells** |  |  |  |  |
| InitSTEM_LambdaSurface | $\lambda_{0s,stem}$ | 2.0 | — | How strongly stem cell regulates its surface area towards a desired size |
| InitSTEM_TargetSurface | $S_{0target,stem}$ | 18.0 | (49) | The ideal surface area each stem cell tries to maintain. Defined to constraint cell shape |
| InitSTEM_LambdaVolume | $\lambda_{0v,stem}$ | 2.0 | — | How strongly a stem cell regulates its volume towards a desired size. |
| InitSTEM_TargetVolume | $V_{0target,stem}$ | 25.0 | (83) | The ideal volume each stem cell tries to maintain. Reference 10 -30 $\mu\text{m}$ |
| DensitySTEM_HalfMaxValue | $k_{mdensity,stem}$ | 125.0 | Fitted (S1.5) | Cell density of stem cells half-max growth response |
| EGF_STEM_HalfMaxValue | $k_{mEGF,stem}$ | 3.5 | Fitted (S1.5) | EGF concentration stem cells achieve half-max growth response |
| InitSTEM_LambdaChemo | $\lambda_{chemoMbias,stem}$ | 100.0 | — | How strongly stem cells move toward areas of higher chemoattract. |
| **Basal Cells** |  |  |  |  |
| InitBASAL_LambdaSurface | $\lambda_{0s,basal}$ | 2.0 | — | How strongly basal cell regulate its surface area towards a desired size |
| InitBASAL_TargetSurface | $S_{0target,basal}$ | 20.0 | (49) | The ideal surface area each basal cell tries to maintain. Defined to constraint cell shape |
| InitBASAL_LambdaVolume | $\lambda_{0v,basal}$ | 2.0 | — | How strongly a basal cell regulates its volume towards a desired size |
| InitBASAL_TargetVolume | $V_{0target,basal}$ | 25.0 | (83) | The ideal volume each basal cell tries to maintain. Reference 10 -30 $\mu\text{m}$ |
| DensityBASAL_HalfMaxValue | $k_{mdensity,basal}$ | 125.0 | Fitted (S1.5) | Cell density of basal cells half-max growth response |
| EGF_BASAL_HalfMaxValue | $k_{mEGF,basal}$ | 7.0 | Fitted (S1.5) | EGF concentration basal cells achieve half-max growth response |
| InitBASAL_LambdaChemo | $\lambda_{chemoMbias,basal}$ | 1000.0 | — | How strongly basal cells move to areas with more chemoattractant |
| **Wing Cells** |  |  |  |  |
| InitWING_LambdaSurface | $\lambda_{0s,wing}$ | 5.0 | — | How strongly wing cell regulates its surface area towards a desired size |
| InitWING_TargetSurface | $S_{0target,wing}$ | 25.0 | (49) | The ideal surface area each wing cell tries to maintain. Defined to constraint cell shape |
| InitWING_LambdaVolume | $\lambda_{0v,wing}$ | 2.0 | — | How strongly a wing cell regulates its volume towards a desired size |
| InitWING_TargetVolume | $V_{0target,wing}$ | 25.0 | (83) | The ideal volume each wing cell tries to maintain. Reference 10 -30 $\mu\text{m}$ |
| InitWING_EGFLambdaChemo | $\lambda_{chemoEGF,wing}$ | 20.0 | (102) | How strongly wing cells move toward regions with higher EGF |
| **Superficial Cells** |  |  |  |  |
| InitSUPER_LambdaSurface | $\lambda_{0s,super}$ | 5.0 | — | How strongly superficial cells regulate surface area to desired size |
| InitSUPER_TargetSurface | $S_{0target,super}$ | 25.0 | (49) | The ideal surface area each superficial cell tries to maintain. Defined to constraint cell shape |
| InitSUPER_LambdaVolume | $\lambda_{0v,super}$ | 5.0 | — | How strongly superficial cells regulate its volume to desired size |
| InitSUPER_TargetVolume | $V_{0target,super}$ | 25.0 | (83) | Ideal volume superficial cell tries to maintain. Reference 10 -30 $\mu\text{m}$ |

**Table S8.2.** Initial parameters (Chemical Fields [EGF, SLS, Movement Bias])

| Parameter | Symbol | Simulation Value | Literature Ref. | Description |
| --- | --- | --- | --- | --- |
| **SLS Diffusion** |  |  |  |  |
| SLS_STEMDiffCoef | $D_{chemstem}$ | 5.0 | — | How quickly the chemical spreads inside stem cells |
| SLS_BASALDiffCoef | $D_{chembasal}$ | 5.0 | — | How quickly the chemical spreads inside basal cells |
| SLS_WINGDiffCoef | $D_{chemwing}$ | 5.0 | — | How quickly the chemical spreads inside wing cells |
| SLS_SUPERDiffCoef | $D_{chemsuper}$ | 5.0 | — | How quickly the chemical spreads inside superficial cells |
| SLS_MEMBDiffCoef | $D_{chemmemb}$ | 5.0 | — | How quickly the chemical spreads in the Bowman's membrane (periphery) |
| SLS_LIMBDiffCoef | $D_{chemlimb}$ | 5.0 | — | How quickly chemical spreads in the Bowman's membrane (limbal region) |
| SLS_TEARDiffCoef | $D_{chemtear}$ | 5.0 | — | How quickly the chemical spreads in the tear layer |
| SLS Global Decay | $k_{dchem}$ | 0.0 | — | For the model the only breakdown and inactivation of the chemical occurs by cell death where the amount inside the cell gets inactive as cells die and washed by the tear |
| **EGF Diffusion / Uptake** |  |  |  |  |
| EGF_SUPERDiffCoef | $D_{EGFsuper}$ | 20.0 | (50,55) | How quickly EGF diffuses (spreads) through superficial cells. The references show a limitation of space for EGF to diffuse through the sides of the cells and it was shown that EGF applied in intact corneas do not proliferate as wounded ones |
| EGF_FieldUptakeBASAL | $\mu_{EGFbasal}$ | 0.0 | — | How quickly basal cells consume EGF from their surroundings |
| EGF_FieldUptakeSTEM | $\mu_{EGFstem}$ | 0.0 | — | How quickly stem cells consume EGF from their surroundings |
| EGF_FieldUptakeSuper | $\mu_{EGFsuper}$ | 0.0 | — | How quickly superficial cells consume EGF |
| EGF_FieldUptakeWing | $\mu_{EGFwing}$ | 0.0 | — | How quickly wing cells consume EGF |
| EGF_ScreteAmount | $\varphi_{EGFtear}$ | 1.0 | (35,54) | The constant amount of EGF introduced into the system from tears. The value for secretion was normalized to have the max value of 1 throughout the simulated space. |
| EGF_GlobalDecay | $k_{dEGF}$ | 0.5 | (88) | The overall rate at which EGF naturally breaks down (decays) over time. The values of EGF half-life from the reference was 78 min ± 36 min transformed into $\approx 0.1$ per MCS, value used is higher to account for other things like uptake, evaporation and sequestration. |
| **Movement Bias Field** |  |  |  |  |
| MovementBiasScreteAmount | $\varphi_{Mbiasmemb, limb}$ | 1.0 | — | The constant amount of chemoattractant secreted by the boundary (Bowman's membrane) to guide cell movement. |
| MovementBiasUptake | $\mu_{Mbiasbasal, stem}$ | 1.0 | — | How quickly basal cells absorb (take up) this chemoattractant |

**Table S8.3.** Initial parameters (Injury Setup and Focal Point Plasticity Links)

| Parameter | Symbol | Simulation Value | Literature Ref. | Description |
| --- | --- | --- | --- | --- |
| SimTime | $t_{final}$ | 50,400.0 | — | How long the simulation runs before it stops (in MCS) |
| **Injury Setup** |  |  |  |  |
| InjuryTime | $t_{injury}$ | 7200.0 | — | The simulation time at which the injury event occurs |
| SLS_X_Center | $x_{centerchem}$ | 100 | — | The x-coordinate of the center of the chemical source |
| SLS_Y_Center | $y_{centerchem}$ | 75 | — | The y-coordinate of the center of the chemical source |
| SLS_Concentration | $c_{chem}$ | [750 a.u.(slight),<br>1500 a.u.(mild),<br>2500 a.u.(mod.)] | (3) | The initial strength or concentration of the chemical introduced. This parameter was chosen based on the descriptions of different classifications following the depth of injury model |
| InjuryX_Center | $x_{centerabla}$ | 150 | — | The x-coordinate of the ablation injury center |
| InjuryY_Center | $y_{centerabla}$ | 60 | — | The y-coordinate of the ablation injury center |
| InjuryRadius | $\pi_{abla}$ | 25 | — | How far from the center cells are affected by the ablation (a radius) |
| **FPP Link Setup** |  |  |  |  |
| LINKWALL_lambda_distance | $\lambda_{0link,wall}$ | 50 | — | How strongly the link between superficial cells and simulation boundary maintains a set distance |
| LINKWALL_target_distance | $L_{0targetlink,wall}$ | 3 | — | Desired length of the link between superficial cells and the boundary |
| LINKWALL_max_distance | $L_{mlink,wall}$ | 1000 | — | Max length link between superficial cells vs boundary can stretch before it breaks |
| LINKSUPER_lambda_distance | $\lambda_{0link,super}$ | 50 | — | How strongly the link between two superficial cells maintains a set distance |
| LINKSUPER_target_distance | $L_{0targetlink,super}$ | 3 | — | The desired resting length of the link between superficial cells |
| LINKSUPER_max_distance | $L_{mlink,super}$ | 1000 | — | Max length the link between superficial cells can stretch before it breaks |

**Table S8.4.**
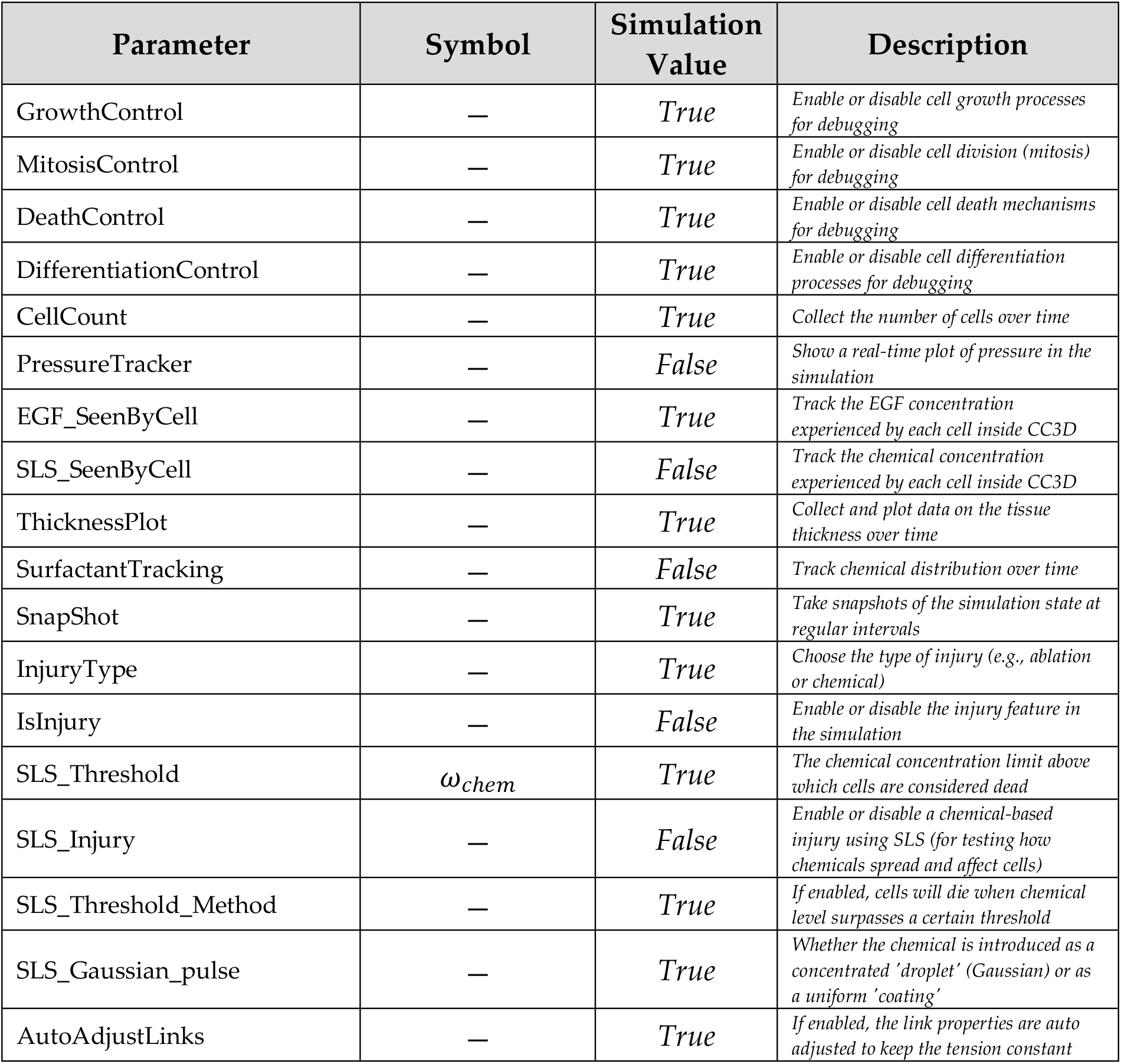
Initial parameters (Simulation/Logging Controls)

## References

1. Clippinger AJ, Raabe HA, Allen DG, Choksi NY, van der Zalm AJ, Kleinstreuer NC, et al. Human-relevant approaches to assess eye corrosion/irritation potential of agrochemical formulations. Cutan Ocul Toxicol. 2021 Apr 3;40(2):145–67.

2. Maurer JK, Parker RD, Jester JV. Extent of Initial Corneal Injury as the Mechanistic Basis for Ocular Irritation: Key Findings and Recommendations for the Development of Alternative Assays. Regul Toxicol Pharmacol. 2002 Aug;36(1):106–17.

3. Scott L, Eskes C, Hoffmann S, Adriaens E, Alepée N, Bufo M, et al. A proposed eye irritation testing strategy to reduce and replace in vivo studies using Bottom–Up and Top– Down approaches. Toxicol In Vitro. 2010 Feb 1;24(1):1–9.

4. Eiferman RA, Schultz GS. Growth Factors and Corneal Epithelium. Cornea. 1987;6(2):149.

5. Kirazowa T, Kinoshira S, Fujira j K, Araki K, Waranabe H, Ohashi Y, et al. The mechanism of accelerated corneal epithelial healing by human epidermal growth factor. Invest Ophthalmol Vis Sci [Internet]. 1990 Sep 1 [cited 2024 Sep 30]; Available from: https://www.semanticscholar.org/paper/The-mechanism-of-accelerated-corneal-epithelial-by-Kirazowa-Kinoshira/7aab78b954bc159fd618606581d5ed55b774062d

6. Maurer JK, Li HF, Petroll WM, Parker RD, Cavanagh HD, Jester JV. Confocal Microscopic Characterization of Initial Corneal Changes of Surfactant-Induced Eye Irritation in the Rabbit. Toxicol Appl Pharmacol. 1997 Apr;143(2):291–300.

7. Zhu J, Lan X, Mo K, Zhang W, Huang Y, Tan J, et al. Deficiency of SECTM1 impairs corneal wound healing in aging. Aging Cell. 2024 Oct;23(10):e14247.

8. Gautheron P, Giroux J, Cottin M, Audegond L, Morilla A, Mayordomo-Blanco L, et al. Interlaboratory assessment of the bovine corneal opacity and permeability (BCOP) assay. Toxicol In Vitro. 1994 Jun 1;8(3):381–92.

9. Schutte K, Prinsen MK, McNamee PM, Roggeband R. The isolated chicken eye test as a suitable in vitro method for determining the eye irritation potential of household cleaning products. Regul Toxicol Pharmacol RTP. 2009 Aug;54(3):272–81.

10. Furukawa M, Sakakibara T, Itoh K, Kawamura K, Matsuura M, Kojima H. Suggestion of the updated IVIS cut-off values for identifying non-ocular irritants in the bovine corneal opacity and permeability (BCOP) assay. Toxicol In Vitro. 2017 Dec 1;45:19–24.

11. Kolle SN, Van Cott A, van Ravenzwaay B, Landsiedel R. Lacking applicability of in vitro eye irritation methods to identify seriously eye irritating agrochemical formulations: Results of bovine cornea opacity and permeability assay, isolated chicken eye test and the EpiOcular^TM^ ET-50 method to classify according to UN GHS. Regul Toxicol Pharmacol. 2017 Apr 1;85:33–47.

12. Jung KM, Lee SH, Ryu YH, Jang WH, Jung HS, Han JH, et al. A new 3D reconstituted human corneal epithelium model as an alternative method for the eye irritation test. Toxicol Vitro Int J Publ Assoc BIBRA. 2011 Feb;25(1):403–10.

13. Sakaguchi H, Ota N, Omori T, Kuwahara H, Sozu T, Takagi Y, et al. Validation study of the Short Time Exposure (STE) test to assess the eye irritation potential of chemicals. Toxicol Vitro Int J Publ Assoc BIBRA. 2011 Jun;25(4):796–809.

14. Swat MH, Thomas GL, Belmonte JM, Shirinifard A, Hmeljak D, Glazier JA. Multi-scale modeling of tissues using CompuCell3D. Methods Cell Biol. 2012;110:325–66.

15. Knudsen TB, Daston GP. 12.23 - Virtual Tissues and Developmental Systems Biology. In: McQueen CA, editor. Comprehensive Toxicology (Second Edition) [Internet]. Oxford: Elsevier; 2010 [cited 2024 May 21]. p. 347–58. Available from: https://www.sciencedirect.com/science/article/pii/B9780080468846015323

16. Shah I, Wambaugh J. Virtual Tissues in Toxicology. J Toxicol Environ Health Part B. 2010 Jun 17;13(2–4):314–28.

17. Clegg LE, Mac Gabhann F. Molecular Mechanism Matters: Benefits of mechanistic computational models for drug development. Pharmacol Res. 2015 Sep;99:149–54.

18. Geris L, editor. Computational Modeling in Tissue Engineering [Internet]. Berlin, Heidelberg: Springer; 2013 [cited 2025 Mar 25]. (Studies in Mechanobiology, Tissue Engineering and Biomaterials; vol. 10). Available from: https://link.springer.com/10.1007/978-3-642-32563-2

19. Hassaniardekani H, Varner V, Petroll M. Computational modeling of the impact of corneal surgical procedures on regional stress distribution within the stroma. Invest Ophthalmol Vis Sci. 2021 Jun 21;62(8):788.

20. Joo J, Kim B, Chun H. Application of computational modeling to improve cornea transplant surgery. J Korean Phys Soc. 2021 Nov 1;79(9):874–81.

21. McDonald TO, Cheng YC, Graser C, Nicol PB, Temko D, Michor F. Computational approaches to modelling and optimizing cancer treatment. Nat Rev Bioeng. 2023 Oct;1(10):695–711.

22. Pak J, Chen ZJ, Sun K, Przekwas A, Walenga R, Fan J. Computational Modeling of Drug Transport Across the In Vitro Cornea. Comput Biol Med. 2018 Jan 1;92:139–46.

23. Nejad TM, Foster C, Gongal D. Finite element modelling of cornea mechanics: a review. Arq Bras Oftalmol [Internet]. 2014 [cited 2024 Nov 10];77(1). Available from: http://www.gnresearch.org/doi/10.5935/0004-2749.20140016

24. Pang G, Wang C, Wang X, Li X, Meng Q. A review of human cornea finite element modeling: geometry modeling, constitutive modeling, and outlooks. Front Bioeng Biotechnol. 2024 Oct 15;12:1455027.

25. Mascolini MV, Toniolo I, Carniel EL, Fontanella CG. Ex vivo, in vivo and in silico studies of corneal biomechanics: a systematic review. Phys Eng Sci Med. 2024 Jun;47(2):403–41.

26. Loerakker S, Ristori T. Computational modeling for cardiovascular tissue engineering: the importance of including cell behavior in growth and remodeling algorithms. Curr Opin Biomed Eng. 2020 Sep;15:1–9.

27. Alshammari FSL. Mathematical modelling of human corneal oxygenation and cell kinetics in corneal epithelial wound healing [Internet] [phd]. Queensland University of Technology; 2016 [cited 2024 Nov 14]. Available from: https://eprints.qut.edu.au/92835/

28. Gaffney EA, Maini PK, Sherratt JA, Tuft S. The Mathematical Modelling of Cell Kinetics in Corneal Epithelial Wound Healing. J Theor Biol. 1999 Mar;197(1):15–40.

29. Moraki E, Grima R, Painter KJ. A stochastic model of corneal epithelium maintenance and recovery following perturbation.

30. Strinkovsky L, Havkin E, Shalom-Feuerstein R, Savir Y. Spatial correlations constrain cellular lifespan and pattern formation in corneal epithelium homeostasis. eLife. 2021 Jan 12;10:e56404.

31. Sridhar MS. Anatomy of cornea and ocular surface. Indian J Ophthalmol. 2018 Feb;66(2):190–4.

32. Chang AY, Purt B. Biochemistry, Tear Film. In: StatPearls [Internet]. Treasure Island (FL): StatPearls Publishing; 2025 [cited 2025 Mar 25]. Available from: http://www.ncbi.nlm.nih.gov/books/NBK572136/

33. Harrell CR, Feulner L, Djonov V, Pavlovic D, Volarevic V. The Molecular Mechanisms Responsible for Tear Hyperosmolarity-Induced Pathological Changes in the Eyes of Dry Eye Disease Patients. Cells. 2023 Dec 1;12(23):2755.

34. Peters E, Colby K. Chapter 3 The Tear Film. In 2009 [cited 2024 Sep 30]. Available from: https://www.semanticscholar.org/paper/Chapter-3-The-Tear-Film-Peters-Colby/62211b36868778b8547e5bb7df218d80d677c802

35. Rao K, Farley WJ, Pflugfelder SC. Association between High Tear Epidermal Growth Factor Levels and Corneal Subepithelial Fibrosis in Dry Eye Conditions. Invest Ophthalmol Vis Sci. 2010 Feb 1;51(2):844–9.

36. McKay TB, Schlötzer-Schrehardt U, Pal-Ghosh S, Stepp MA. Integrin: Basement membrane adhesion by corneal epithelial and endothelial cells. Exp Eye Res. 2020 Sep;198:108138.

37. Streuli CH, Schmidhauser C, Bailey N, Yurchenco P, Skubitz AP, Roskelley C, et al. Laminin mediates tissue-specific gene expression in mammary epithelia. J Cell Biol. 1995 May;129(3):591–603.

38. Chaurasia SS, Kaur H, De Medeiros FW, Smith SD, Wilson SE. Reprint of “Dynamics of the expression of intermediate filaments vimentin and desmin during myofibroblast differentiation after corneal injury.” Exp Eye Res. 2009 Oct;89(4):590–6.

39. Hynes RO. Extracellular matrix: not just pretty fibrils. Science. 2009 Nov 27;326(5957):1216–9.

40. Ingber DE. Cellular mechanotransduction: putting all the pieces together again. FASEB J Off Publ Fed Am Soc Exp Biol. 2006 May;20(7):811–27.

41. Nakamura K. Intact corneal epithelium is essential for the prevention of stromal haze after laser assisted in situ keratomileusis. Br J Ophthalmol. 2001 Feb 1;85(2):209–13.

42. Singh V, Santhiago MR, Barbosa FL, Agrawal V, Singh N, Ambati BK, et al. Effect of TGFβ and PDGF-B blockade on corneal myofibroblast development in mice. Exp Eye Res. 2011 Dec;93(6):810–7.

43. Torricelli AAM, Singh V, Santhiago MR, Wilson SE. The Corneal Epithelial Basement Membrane: Structure, Function, and Disease. Invest Ophthalmol Vis Sci. 2013 Sep;54(9):6390–400.

44. Torricelli AAM, Santhanam A, Wu J, Singh V, Wilson SE. The corneal fibrosis response to epithelial-stromal injury. Exp Eye Res. 2016 Jan;142:110–8.

45. Schlötzer-Schrehardt U, Dietrich T, Saito K, Sorokin L, Sasaki T, Paulsson M, et al. Characterization of extracellular matrix components in the limbal epithelial stem cell compartment. Exp Eye Res. 2007 Dec;85(6):845–60.

46. Dorà NJ, Hill RE, Collinson JM, West JD. Lineage tracing in the adult mouse corneal epithelium supports the limbal epithelial stem cell hypothesis with intermittent periods of stem cell quiescence. Stem Cell Res. 2015 Nov;15(3):665–77.

47. Dziasko MA, Daniels JT. Anatomical Features and Cell-Cell Interactions in the Human Limbal Epithelial Stem Cell Niche. Ocul Surf. 2016 Jul;14(3):322–30.

48. Beebe DC, Masters BR. Cell lineage and the differentiation of corneal epithelial cells. Invest Ophthalmol Vis Sci. 1996 Aug;37(9):1815–25.

49. Sterenczak KA, Winter K, Sperlich K, Stahnke T, Linke S, Farrokhi S, et al. Morphological characterization of the human corneal epithelium by in vivo confocal laser scanning microscopy. Quant Imaging Med Surg. 2021 May;11(5):1737–50.

50. Van Itallie CM, Holmes J, Bridges A, Gookin JL, Coccaro MR, Proctor W, et al. The density of small tight junction pores varies among cell types and is increased by expression of claudin-2. J Cell Sci. 2008 Feb 1;121(3):298–305.

51. Ban Y, Dota A, Cooper LJ, Fullwood NJ, Nakamura T, Tsuzuki M, et al. Tight junction-related protein expression and distribution in human corneal epithelium. Exp Eye Res. 2003 Jun 1;76(6):663–9.

52. Estil S, Primo EJ, Wilson G. Apoptosis in Shed Human Corneal Cells. Invest Ophthalmol Vis Sci. 2000 Oct 1;41(11):3360–4.

53. Yoon JJ, Ismail S, Sherwin T. Limbal stem cells: Central concepts of corneal epithelial homeostasis. World J Stem Cells. 2014 Sep 26;6(4):391–403.

54. Peterson JL, Ceresa BP. Epidermal Growth Factor Receptor Expression in the Corneal Epithelium. Cells. 2021 Sep 13;10(9):2409.

55. Savage CR, Cohen S. Proliferation of corneal epithelium induced by epidermal growth factor. Exp Eye Res. 1973 Mar 1;15(3):361–6.

56. Wee P, Wang Z. Epidermal Growth Factor Receptor Cell Proliferation Signaling Pathways. Cancers. 2017 May 17;9(5):52.

57. Gouveia RM, Lepert G, Gupta S, Mohan RR, Paterson C, Connon CJ. Assessment of corneal substrate biomechanics and its effect on epithelial stem cell maintenance and differentiation. Nat Commun. 2019 Apr 3;10(1):1496.

58. Ouyang H, Xue Y, Lin Y, Zhang X, Xi L, Patel S, et al. WNT7A and PAX6 define corneal epithelium homeostasis and pathogenesis. Nature. 2014 Jul;511(7509):358–61.

59. Gandhi S, Jain S. The Anatomy and Physiology of Cornea. In: Cortina MS, de la Cruz J, editors. Keratoprostheses and Artificial Corneas: Fundamentals and Surgical Applications [Internet]. Berlin, Heidelberg: Springer; 2015 [cited 2024 Jul 18]. p. 19–25. Available from: 10.1007/978-3-642-55179-6_3

60. Halkiadakis I, Vernikou A, Tzimis V, Markopoulos I, Popeskou K, Konstadinidou V. Assessment of Corneal Epithelium Thickness in Glaucomatous Patients Undergoing Medical Treatment. J Glaucoma. 2021 Jan;30(1):44.

61. Kimura K, Kawano S, Mori T, Inoue J, Hadachi H, Saito T, et al. Quantitative Analysis of the Effects of Extracellular Matrix Proteins on Membrane Dynamics Associated with Corneal Epithelial Cell Motility. Invest Ophthalmol Vis Sci. 2010 Sep 1;51(9):4492–9.

62. Kariya Y, Miyazaki K. The basement membrane protein laminin-5 acts as a soluble cell motility factor. Exp Cell Res. 2004 Jul 15;297(2):508–20.

63. Park M, Richardson A, Pandzic E, Lobo EP, Whan R, Watson SL, et al. Visualizing the Contribution of Keratin-14+ Limbal Epithelial Precursors in Corneal Wound Healing. Stem Cell Rep. 2019 Jan;12(1):14–28.

64. Lobo EP, Delic NC, Richardson A, Raviraj V, Halliday GM, Di Girolamo N, et al. Self-organized centripetal movement of corneal epithelium in the absence of external cues. Nat Commun. 2016 Aug 8;7:12388.

65. Richardson A, Lobo EP, Delic NC, Myerscough MR, Lyons JG, Wakefield D, et al. Keratin-14-Positive Precursor Cells Spawn a Population of Migratory Corneal Epithelia that Maintain Tissue Mass throughout Life. Stem Cell Rep. 2017 Oct 10;9(4):1081–96.

66. Wilson SE, Liang Q, Kim WJ. Lacrimal Gland HGF, KGF, and EGF mRNA Levels Increase after Corneal Epithelial Wounding. Invest Ophthalmol Vis Sci. 1999 Sep 1;40(10):2185–90.

67. Ohashi Y, Motokura M, Kinoshita Y, Mano T, Watanabe H, Kinoshita S, et al. Presence of epidermal growth factor in human tears. Invest Ophthalmol Vis Sci. 1989 Aug;30(8):1879– 82.

68. Iglesia DDSta, Stepp MA. Disruption of the Basement Membrane after Corneal Débridement. Invest Ophthalmol Vis Sci. 2000 Apr 1;41(5):1045–53.

69. Wilson SE. Fibrosis Is a Basement Membrane-Related Disease in the Cornea: Injury and Defective Regeneration of Basement Membranes May Underlie Fibrosis in Other Organs. Cells. 2022 Jan;11(2):309.

70. Akowuah PK, De La Cruz A, Smith CW, Rumbaut RE, Burns AR. An Epithelial Abrasion Model for Studying Corneal Wound Healing. J Vis Exp JoVE. 2021 Dec 29;(178):10.3791/63112.

71. Domingo E, Moshirfar M, Zeppieri M, Zabbo CP. Corneal Abrasion. In: StatPearls [Internet]. Treasure Island (FL): StatPearls Publishing; 2025 [cited 2025 Mar 28]. Available from: http://www.ncbi.nlm.nih.gov/books/NBK532960/

72. Ramamurthi S, Rahman MQ, Dutton GN, Ramaesh K. Pathogenesis, clinical features and management of recurrent corneal erosions. Eye. 2006 Jun;20(6):635–44.

73. Sridhar MS, Rapuano CJ, Cosar CB, Cohen EJ, Laibson PR. Phototherapeutic keratectomy versus diamond burr polishing of Bowman’s membrane in the treatment of recurrent corneal erosions associated with anterior basement membrane dystrophy. Ophthalmology. 2002 Apr 1;109(4):674–9.

74. Espana EM, Birk DE. Composition, Structure and Function of the Corneal Stroma. Exp Eye Res. 2020 Sep;198:108137.

75. Kamil S, Mohan RR. Corneal stromal wound healing: Major regulators and therapeutic targets. Ocul Surf. 2021 Jan;19:290–306.

76. Maurice DM. The structure and transparency of the cornea. J Physiol. 1957 Apr 30;136(2):263-286.1.

77. Pavelka M, Roth J. Descemet’s Membrane. In: Pavelka M, Roth J, editors. Functional Ultrastructure: Atlas of Tissue Biology and Pathology [Internet]. Vienna: Springer; 2010 [cited 2024 Jul 18]. p. 184–5. Available from: 10.1007/978-3-211-99390-3_97

78. Santerre K, Xu I, Thériault M, Proulx S. In Vitro Expansion of Corneal Endothelial Cells for Transplantation. Methods Mol Biol Clifton NJ. 2020;2145:17–27.

79. Niederkorn JY. Corneal transplantation and immune privilege. Int Rev Immunol. 2013 Feb;32(1):57–67.

80. Streilein JW. Ocular immune privilege: the eye takes a dim but practical view of immunity and inflammation. J Leukoc Biol. 2003 Aug;74(2):179–85.

81. Graner F, Glazier JA. Simulation of biological cell sorting using a two-dimensional extended Potts model. Phys Rev Lett. 1992 Sep 28;69(13):2013–6.

82. Montagud A, Ponce-de-Leon M, Valencia A. Systems biology at the giga-scale: Large multiscale models of complex, heterogeneous multicellular systems. Curr Opin Syst Biol. 2021 Dec;28:100385.

83. De Paiva CS, Pflugfelder SC, Li DQ. Cell Size Correlates with Phenotype and Proliferative Capacity in Human Corneal Epithelial Cells. Stem Cells. 2006 Feb 1;24(2):368–75.

84. Abu-Romman A, Scholand KK, Govindarajan G, Yu Z, Pal-Ghosh S, Stepp MA, et al. Age-Related Differences in the Mouse Corneal Epithelial Transcriptome and Their Impact on Corneal Wound Healing. Invest Ophthalmol Vis Sci. 2024 May 13;65(5):21.

85. Shadmani A, Wu AY. Navigating the path to corneal healing success and challenges: a comprehensive overview. Eye. 2025 Feb 12;1–9.

86. Zhang Y, Do KK, Wang F, Lu X, Liu JY, Li C, et al. Zeb1 facilitates corneal epithelial wound healing by maintaining corneal epithelial cell viability and mobility. Commun Biol. 2023 Apr 20;6(1):1–12.

87. Kuo CY, Eranki A, Placone JK, Rhodes KR, Aranda-Espinoza H, Fernandes R, et al. Development of a 3D Printed, Bioengineered Placenta Model to Evaluate the Role of Trophoblast Migration in Preeclampsia. ACS Biomater Sci Eng. 2016 Oct 10;2(10):1817– 26.

88. Chan KY, Lindquist TD, Edenfield MJ, Nicolson MA, Banks AR. Pharmacokinetic study of recombinant human epidermal growth factor in the anterior eye. Invest Ophthalmol Vis Sci. 1991 Dec;32(13):3209–15.

89. Ljubimov AV, Saghizadeh M. Progress in corneal wound healing. Prog Retin Eye Res. 2015 Nov;49:17–45.

90. Lu L, Reinach PS, Kao WW. Corneal epithelial wound healing. Exp Biol Med Maywood NJ. 2001 Jul;226(7):653–64.

91. Vaidyanathan U, Hopping GC, Liu HY, Somani AN, Ronquillo YC, Hoopes PC, et al. Persistent Corneal Epithelial Defects: A Review Article. Med Hypothesis Discov Innov Ophthalmol. 2019;8(3):163–76.

92. Shahid SM, Harrison N. Corneal abrasion: assessment and management. InnovAiT. 2013 Sep 1;6(9):551–4.

93. Ahmed F, House RJ, Feldman BH. Corneal Abrasions and Corneal Foreign Bodies. Prim Care. 2015 Sep;42(3):363–75.

94. Paley GL, Wagoner MD, Afshari NA, Pineda R, Huang AJW, Kenyon KR. Corneal Wound Healing, Recurrent Corneal Erosions, and Persistent Epithelial Defects. In: Albert DM, Miller JW, Azar DT, Young LH, editors. Albert and Jakobiec’s Principles and Practice of Ophthalmology [Internet]. Cham: Springer International Publishing; 2022 [cited 2024 Nov 18]. p. 331–60. Available from: 10.1007/978-3-030-42634-7_212

95. Ebihara N, Mizushima H, Miyazaki K, Watanabe Y, Ikawa S, Nakayasu K, et al. The Functions of Exogenous and Endogenous Laminin-5 on Corneal Epithelial Cells. Exp Eye Res. 2000 Jul;71(1):69–79.

96. Jadczyk-Sorek K, Garczorz W, Bubała-Stachowicz B, Francuz T, Mrukwa-Kominek E. Matrix Metalloproteinases and the Pathogenesis of Recurrent Corneal Erosions and Epithelial Basement Membrane Dystrophy. Biology. 2023 Sep 21;12(9):1263.

97. Cenedella RJ, Fleschner CR. Kinetics of corneal epithelium turnover in vivo. Studies of lovastatin. Invest Ophthalmol Vis Sci. 1990 Oct;31(10):1957–62.

98. Hirst L, Fogle J, Kenyon K, Stark W. Corneal epithelial regeneration and adhesion following acid burns in the rhesus monkey. Invest Ophthalmol Vis Sci [Internet]. 1982 Dec 1 [cited 2024 Sep 30]; Available from: https://www.semanticscholar.org/paper/Corneal-epithelial-regeneration-and-adhesion-acid-Hirst-Fogle/a6430bacc2406fb78059b7620603aab2d262305f

99. Kenyon K, Fogle J, Stone DL, Stark W. Regeneration of corneal epithelial basement membrane following thermal cauterization. Invest Ophthalmol Vis Sci [Internet]. 1977 Apr 1 [cited 2024 Sep 30]; Available from: https://www.semanticscholar.org/paper/Regeneration-of-corneal-epithelial-basement-thermal-Kenyon-Fogle/04d5c4407a2ac9f10dfd295e861f0c13783e6f2d

100. Alghamdi A, Khan MS, Dakhil TA. Understanding Corneal Epithelial Thickness Mapping. Middle East Afr J Ophthalmol. 2023 May 25;29(3):147–55.

101. Thorne RG, Hrabětová S, Nicholson C. Diffusion of Epidermal Growth Factor in Rat Brain Extracellular Space Measured by Integrative Optical Imaging. J Neurophysiol. 2004 Dec;92(6):3471–81.

102. Boisjoly HM, Laplante C, Bernatchez SF, Salesse C, Giasson M, Joly MC. Effects of EGF, IL-1 and their Combination on In Vitro Corneal Epithelial Wound Closure and Cell Chemotaxis. Exp Eye Res. 1993 Sep 1;57(3):293–300.

